# Discovery of a Potent and Selective Inhibitor of Human NLRP3 with a Novel Binding Modality and Mechanism of Action

**DOI:** 10.1101/2024.12.21.629867

**Authors:** Kevin Wilhelmsen, Aditi Deshpande, Sarah Tronnes, Maitriyee Mahanta, Matthew Banicki, Mary Cochran, Samantha Cowdin, Kristen Fortney, George Hartman, Robert Hughes, Rusty Montgomery, Claudia Portillo, Paul Rubin, Yan Wang, Shijun Yan, Barry A Morgan, Assem Duisembekova, Romane Riou, Michael Marleaux, Inga V. Hochheiser, Hannes Buthmann, Dominic Ferber, Wei Wang, Melanie Cranston, Chloe M. McKee, Thea Mawhinney, Emma McKay, Bénédicte F. Py, Matthias Geyer, Rebecca C. Coll

**Affiliations:** BioAge Labs, 1445 S. 50th St. Richmond, CA 94804, USA; HitGen Pharmaceuticals Inc, PO Box 88240, Houston, TX 77288, USA; CIRI, Centre International de Recherche en Infectiologie, Univ Lyon, Inserm, U1111, Université Claude Bernard Lyon 1, CNRS, UMR5308, ENS de Lyon, Lyon, France; Institute of Structural Biology, University of Bonn, Venusberg-Campus 1, 53127, Bonn, Germany; Wellcome-Wolfson Institute for Experimental Medicine, Queen’s University Belfast, Belfast BT9 7BL, UK

**Keywords:** NLRP3, Inflammasome, Inflammation, Inflammaging, MCC950, CRID3, BAL-0028, Small molecule inhibitor

## Abstract

The NLRP3 inflammasome is an intracellular protein complex that causes inflammation via the release of IL-1β and pyroptosis. NLRP3 activation is associated with many age-related inflammatory diseases, and NLRP3 inhibition is a promising therapeutic strategy. We previously performed a DNA encoded library screen to identify novel NLRP3 binding molecules. Herein we describe the characterization of BAL-0028 as a potent and specific inhibitor of NLRP3 signaling. Notably, BAL-0028 is a poor inhibitor of mouse NLRP3 but inhibits human and primate NLRP3 with nanomolar potency. Using cellular and biochemical analyses we demonstrate that BAL-0028 binds to the NLRP3 NACHT domain at a site that is distinct from the MCC950 binding pocket. Using humanized NLRP3 mice we show that a derivative of BAL-0028 inhibits NLRP3 activation *in vivo* in a peritonitis model. Finally, we demonstrate that BAL-0028 inhibits select hyperactive NLRP3 mutations associated with autoinflammatory diseases more potently than does MCC950. BAL-0028 thus represents a new modality for NLRP3 inhibition in inflammatory diseases.

**SUMMARY:** NLRP3 is a target for anti-inflammatory therapies and can be inhibited by the tool compound MCC950. We describe the characterization of a new small molecule inhibitor of NLRP3 BAL-0028 that has a distinct mechanism of action and binding site.

## INTRODUCTION

Inflammation is associated with many age-related diseases including neurodegenerative conditions, cancer, and metabolic and cardiovascular diseases(Franceschi et al., 2018; Ransohoff, 2016). Limiting inflammation may thus be an effective therapeutic strategy to reduce or delay age-related diseases(Figueira et al., 2016). Inflammasomes are protein complexes that have emerged as central mediators of inflammation as they control the production of the pro-inflammatory cytokines interleukin (IL)-1β and IL-18, and lytic inflammatory cell death known as pyroptosis(Broz and Dixit, 2016). Inflammasomes are formed by a pattern recognition receptor (PRR) or sensor molecule that interacts with the adapter molecule apoptosis-associated speck-like protein containing a CARD (ASC). ASC oligomerizes and provides a platform for the autocatalytic activation of the zymogen protease caspase-1. Active caspase-1 cleaves pro-IL-1β and IL-18 into their active secreted forms and mediates pyroptosis via cleavage of gasdermin D (GSDMD)(Fu et al., 2024).

Amongst the known inflammasome sensors, the NACHT, LRR and PYD domains-containing protein 3 (NLRP3) has emerged a key mediator of pathogenic inflammation in many inflammatory diseases. NLRP3 can be activated by a broad range of molecules and processes and is regarded as a sensor of the disruption of cellular homeostasis(Akbal et al., 2022). Disease-related molecules such as β-amyloid, α-synuclein, and monosodium urate crystals have been shown to trigger NLRP3 activation. NLRP3-deficient mice are correspondingly protected in models of Alzheimer’s Disease, Parkinson’s Disease and gout. There is also a group of rare genetic diseases caused by gain-of-function mutations in NLRP3 called Cryopyrin-associated periodic syndromes (CAPS) or NLRP3-associated autoinflammatory diseases (NLRP3-AID). NLRP3 inhibition is thus a promising anti-inflammatory therapeutic strategy(Coll et al., 2022; Mangan et al., 2018).

Currently available therapies to inhibit the inflammasome pathway are limited to antibodies such as Anakinra (IL-1 receptor antagonist) and Canakinumab (anti-IL-1β) (Coll, 2023). However, specific NLRP3 inhibition could have several advantages over these including preventing IL-18 release and pyroptosis, while maintaining inflammation driven by other inflammasomes like NLRP1 and NLRC4 that may be important for host defense(Barnett et al., 2023). In addition, small molecule inhibitors have improved blood-brain barrier crossing, which would be critical for blocking neuroinflammation(Xiong et al., 2021).

Numerous small molecule NLRP3 inhibitors have been described including the sulfonylurea MCC950 (aka CRID3, CP-456,773). MCC950 was developed based on early observations from cell-based screens that identified diarylsulfonylureas such as glyburide could inhibit NLRP3 signaling(Laliberte et al., 2003; Lamkanfi et al., 2009; Perregaux et al., 2001). MCC950 is the most widely used tool molecule to study NLRP3 inhibition in cells and in animal models of disease(Coll et al., 2015; Corcoran et al., 2021). MCC950 is a specific non-covalent inhibitor that blocks the ATPase activity of NLRP3(Coll et al., 2019; Tapia-Abellan et al., 2019; Vande Walle et al., 2019). Its binding site has been resolved at molecular resolution with two reports of NLRP3-MCC950 cryo-EM structures(Hochheiser et al., 2022b; Ohto et al., 2022) and a crystal structure of NLRP3 with an MCC950 analogue(Dekker et al., 2021). Notably the majority of NLRP3 inhibitors that have advanced to early stage clinical trials are sulfonylureas or MCC950-derivatives(Vande Walle and Lamkanfi, 2024).

Several other direct NLRP3 inhibitors with different chemistries and mechanisms of action have been described such as CY-09 which prevents ATP binding, and Tranilast which disrupts NLRP3 oligomerisation(Coll et al., 2022; Huang et al., 2018; Jiang et al., 2017). However, many of these compounds have off-target effects and indeed MCC950 has also been shown to inhibit carbonic anhydrase 2(Coll et al., 2022; Kennedy et al., 2021; Walle et al., 2024). Importantly, in studies of NLRP3-AID mouse models and patient cells it has been observed that MCC950 cannot effectively inhibit NLRP3 activation by certain mutations(Cosson et al., 2024; Vande Walle et al., 2019). There is therefore a need to identify specific NLRP3 inhibitors with novel mechanisms of action that can offer an alternative to MCC950-derived compounds. Herein we describe BAL-0028, a specific inhibitor of NLRP3 with a mechanism that is distinct from MCC950. Remarkably, we find that BAL-0028 potently inhibits primate NLRP3 but has significantly reduced potency for NLRP3 from other mammals, highlighting the importance of drug development efforts focused on human proteins.

## RESULTS

Using a DNA encoded library (DEL) screen with recombinant NLRP3 we identified a series of indazole small molecule NLRP3 binders(Hartman et al., 2024). A high affinity (*K*_D_ range 104-123 nM) lead compound from this study, BAL-0028 (Fig. 1A), was selected for further characterization in cell-based assays(Hartman et al., 2024). We first examined BAL-0028 in NLRP3 signaling assays using THP-1 macrophage-like cells. Pre-treatment with BAL-0028 potently inhibits NLRP3-dependent IL-1β release from THP-1s simulated with LPS and the pore forming toxin nigericin, with an IC_50_ of 57.5 nM. In the same assay the IC_50_ of MCC950 is 14.3 nM which is consistent with previous reports(Clenet et al., 2023; Teske et al., 2024) (Fig. 1B). NLRP3 is activated by many molecules including damage associated molecular patterns (DAMPS) release during sterile inflammation such as ATP, and monosodium urate (MSU) crystals(Kapetanovic et al., 2015; Mangan et al., 2018). BAL-0028 also inhibits IL-1β release triggered by ATP and MSU with IC_50_s in the nanomolar range (Fig. 1C,D). This confirms the ability of BAL-0028 to block NLRP3-dependent signaling by multiple stimuli in THP-1s.

**Figure 1.**
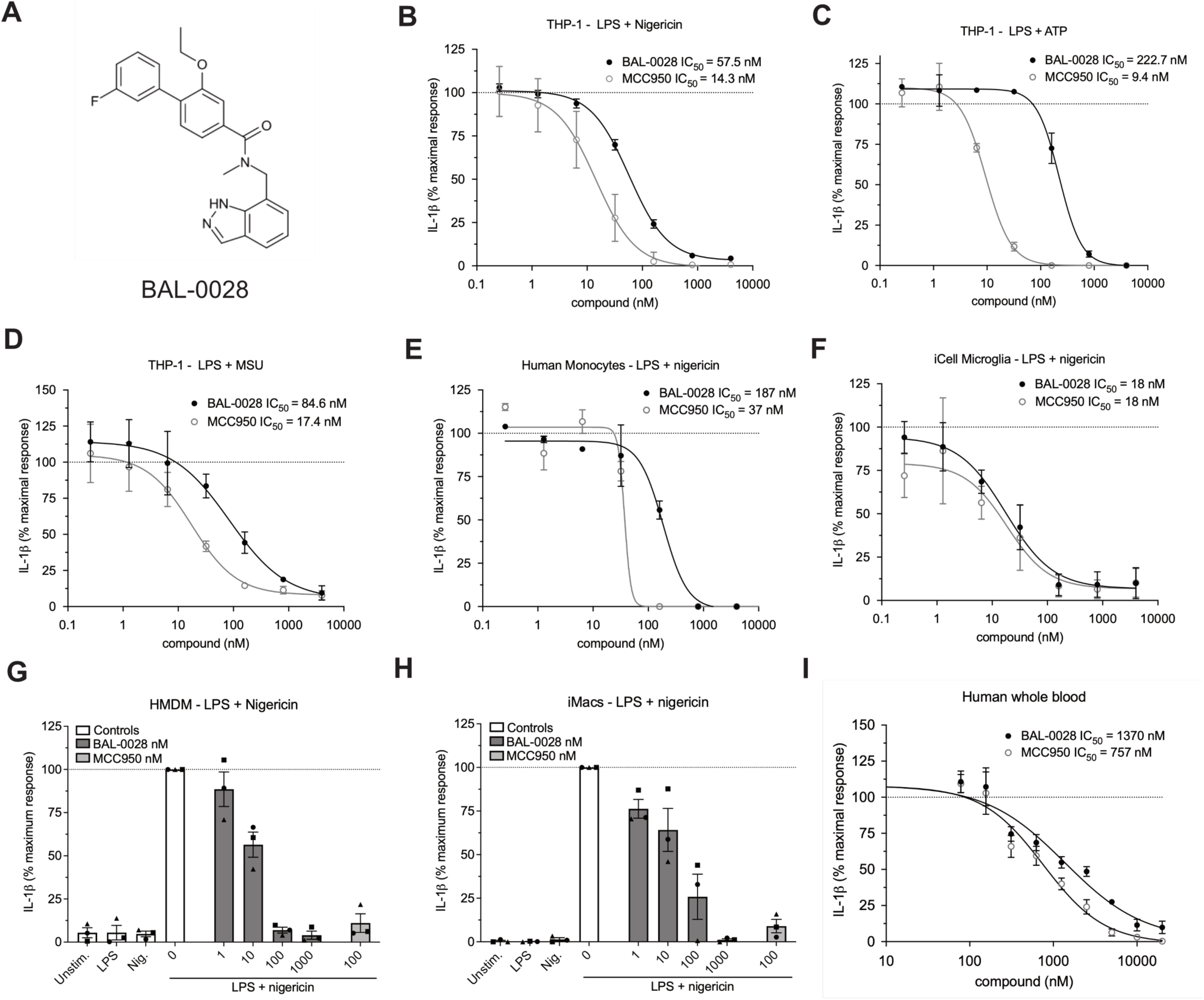
BAL-0028 is a potent inhibitor of NLRP3 signaling in multiple human cell types. (A) Structure of BAL-0028. (B-D) Comparison of BAL-0028 and MCC950 in IL-1β release assays from PMA differentiated THP-1 cells stimulated with LPS and (B) nigericin, (C) ATP, or (D) MSU. (E-I) Comparison of BAL-0028 and MCC950 in IL-1β release assays from LPS and nigericin stimulated human monocytes (E), iCell microglia (F), HMDM (G), iMacs (H), and human whole blood (I). (B-F,I) Graph symbols show average IL-1β values relative to vehicle control +/− S.E.M. from independent experiments performed in triplicate; IC50 curve fitted by non-linear regression analysis. (G-H) Graph symbols show average values relative to vehicle control from independent experiments (indicated by different symbols) performed in triplicate +/− S.E.M. Compounds are shown in nanomolar (nM) concentrations. (B) N=73 for BAL-0028 and N=29 for MCC950, (C-E) N=2 (F,H) N=3, (G) N=3 donors, (I) N=4 donors.

BAL-0028 was further evaluated in a range of more physiologically relevant human cell types. BAL-0028 consistently inhibits nigericin induced IL-1β release in primary monocytes (Fig. 1E) and induced pluripotent stem cell (iPSC)-derived microglia (iCell microglia) (Fig. 1F). The IC_50_ of BAL-0028 is generally higher than MCC950 (Fig. 1B-E), however for iCell microglia the compounds are equipotent (Fig. 1F). Having established the potency of BAL-0028 we tested it in primary human monocyte-derived macrophages (HMDM) and iPSC-derived macrophages (iMacs). BAL-0028 inhibited nigericin induced IL-1β release in HMDM and iMacs in the nanomolar range (Fig. 1G-H). In addition, we examine lactate dehydrogenase (LDH) release as a measure of pyroptotic cell death induced by NLRP3. LDH is dose-dependently inhibited by BAL-0028 in iMacs (Supp. Fig. 1A). Together these data demonstrate that BAL-0028 is a potent inhibitor of NLRP3 signaling in multiple human cell types. Stimulation with LPS is required to induce both transcriptional and posttranslational priming of the NLRP3 inflammasome(McKee and Coll, 2020; O’Keefe et al., 2024). To examine the effects of BAL-0028 on LPS signaling we measured the release of inflammasome-independent inflammatory cytokines TNF and IL-6. BAL-0028 does not reduce TNF secretion from HMDM or iMacs in NLRP3 assays (Supp Fig. 1B). Pre-treatment with BAL-0028 does not block LPS induced TNF secretion, although there was some reduction in IL-6 release at the highest concentration of BAL-0028 (10 mM) (Supp. Fig. 1C-D). These data demonstrate that BAL-0028 does not appreciably interfere with LPS signaling. We also confirmed that BAL-0028 is not cytotoxic as it does not increase LDH release or reduce cell viability in THP-1s (Supp Fig. 1E-F). Lastly, we compared BAL-0028 and MCC950 in a whole blood NLRP3 assay. As expected, there was a decrease in potency of both BAL-0028 and MCC950 due to plasma protein binding, but both compounds effectively inhibited IL-1β release (Fig. 1I). This confirms that BAL-0028 is a potent inhibitor of human NLRP3 in a clinically relevant assay.

We next examined the effects of BAL-0028 on signaling events upstream of IL-1β secretion and pyroptosis. Upon NLRP3 inflammasome activation pro-IL-1β is cleaved into its active p17 form by caspase-1(Broz and Dixit, 2016). Through Western blotting, we confirmed that both BAL-0028 and MCC950 inhibited pro-IL-1β processing and caspase-1 activation as measured by the appearance of the p20 auto-processed form of caspase-1(Boucher et al., 2018) (Fig. 2A). BAL-0028 did not affect the expression of pro-IL-1β or NLRP3, again suggesting NLRP3 priming by LPS is unaffected by BAL-0028 (Fig. 2A). To examine whether NLRP3 inflammasome formation is inhibited by BAL-0028 we measured ASC-speck formation by fluorescence microscopy. In THP-1 cells stably expressing green fluorescent protein (GFP) tagged ASC, nigericin induced ASC speck formation is dose-dependently inhibited by both BAL-0028 and MCC950 (Fig. 2B). We next used HEK293T cells expressing BlueFP-ASC and doxycycline-inducible NLRP3 to monitor ASC specks by flow cytometry. In these cells, nigericin stimulation triggers a significant increase in ASC specks, which is dose-dependently inhibited by both BAL-0028 and MCC950 (Fig. 2C). In addition in iMacs, nigericin-induced ASC speck formation was also potently blocked by BAL-0028 (Fig. 2D,E). These data confirm that BAL-0028 inhibits the formation of the NLRP3 inflammasome.

**Figure 2.**
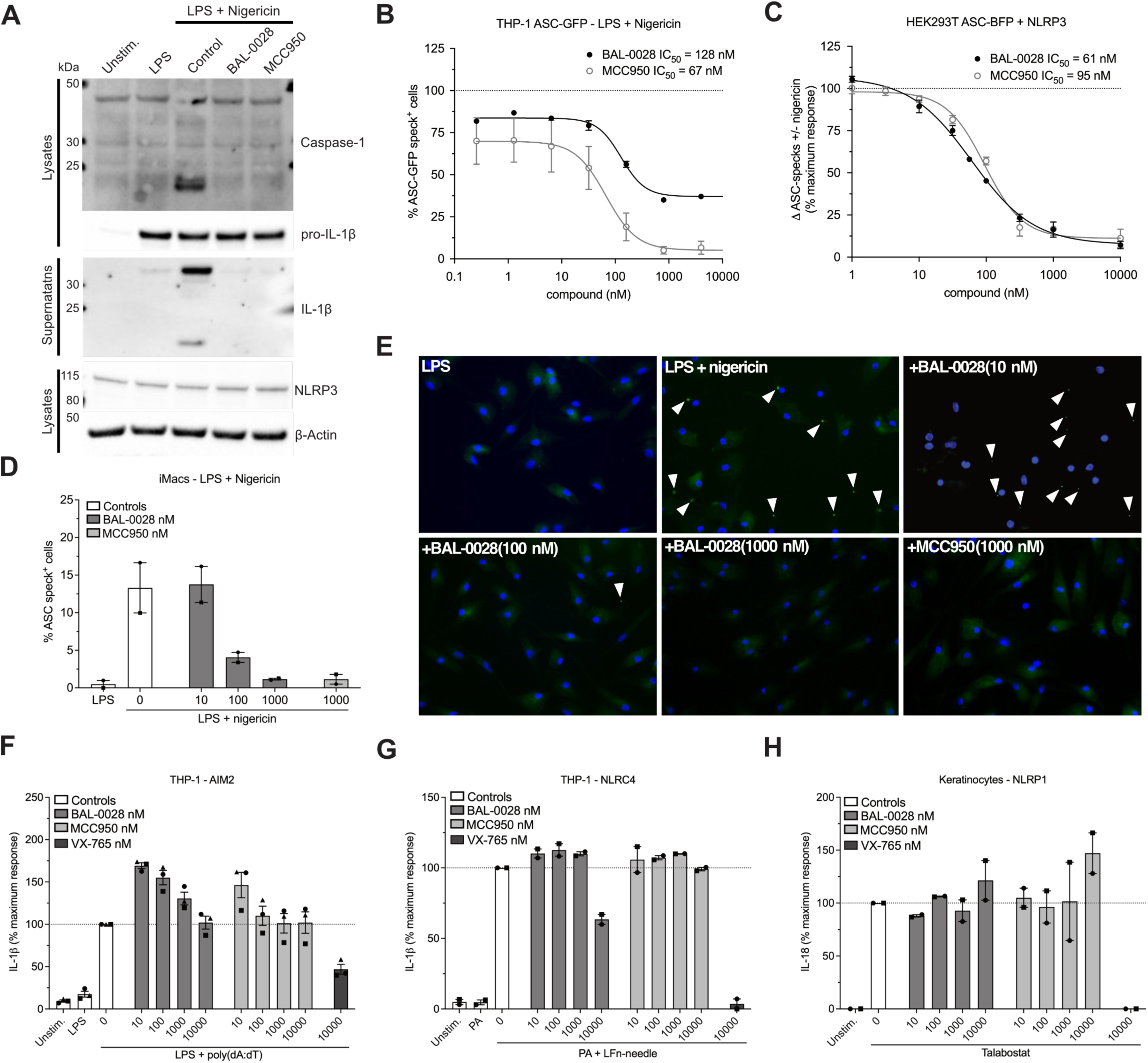
BAL-0028 specifically inhibits NLRP3 inflammasome formation. (A) Western blot for caspase-1 and IL-1β cleavage, and NLRP3 expression from PMA differentiated THP-1 cells stimulated with LPS and nigericin in the presence of BAL-0028 or MCC950 (both 500 nM). (B) Comparison of BAL-0028 and MCC950 effects on ASC speck formation assessed by fluorescence microscopy in PMA differentiated THP-1 ASC-GFP cells stimulated with LPS and nigericin. (C) Comparison of BAL-0028 and MCC950 effects on ASC speck formation assessed by flow cytometry in HEK293T ASC-BFP cells transfected with human NLRP3 and stimulated with nigericin. (D-E) Effect of BAL-0028 and MCC950 on ASC speck formation assessed by fluorescence microscopy using an anti-ASC antibody in iMacs. (F-G) Effects of BAL-0028, MCC950, and VX-765 on IL-1β release from PMA differentiated THP-1 cells stimulated with (F) LPS and transfected with poly(dA:dT) or (G) protective antigen (PA) and Lfn-needle protein. (H) Effects of BAL-0028, MCC950 and VX-765 on IL-18 release from human keratinocytes stimulated with Talabostat. (A) Representative blots from N=2 independent experiments. (B) Average +/− S.E.M % ASC-GFP speck positive cells from N=2 independent experiments performed in triplicate. (C) Average +/− S.E.M change in nigericin induced ASC specks normalized to cells without compound treatment from N=3-4 independent experiments. (D,F-H) Graph symbols show average values relative to vehicle control from independent experiments performed in triplicate (indicated by different symbols) +/− S.E.M. (D,G,H) N=2 and (F) N=3. (E) Representative images from (D).

To confirm the specificity of BAL-0028 for NLRP3, we assessed the effects of BAL-0028 on other inflammasome pathways. Transfection of the synthetic double stranded DNA Poly(dA:dT) in THP-1s induces IL-1β release via the dsDNA sensor AIM2(Fernandes-Alnemri et al., 2009; Hornung et al., 2009). In this AIM2 assay BAL-0028 and MCC950 did not reduce IL-1β release relative to the vehicle control, while the caspase-1 inhibitor VX-765(Wannamaker et al., 2007) attenuated IL-1β release (Fig. 2F). The NAIP/NLRC4 inflammasome senses bacterial infection and can be activated by treatment with a needle protein (LFn-BsaL) and protective antigen (PA)(Matico et al., 2024; Yang et al., 2013). NAIP/NLRC4-dependent IL-1β release was completely blocked by VX-765 but was not inhibited by MCC950 (Fig. 2G). BAL-0028 did not inhibit IL-1β release up to 1 μM but did reduce IL-1β at the highest dose of 10 μM (Fig. 2G). Cell death induced by AIM2 and NLRC4 was not affected by BAL-0028 or MCC950; at 10 μM VX-765 also did not inhibit cell death, as higher concentrations are required to block LDH relative to IL-1β release(Schneider et al., 2017)(Supp. Fig. 2A-B). NLRP1 is highly expressed in human keratinocytes and can be activated by the small molecule dipeptidyl peptidase inhibitor Talabostat (Val-boroPro)(Zhong et al., 2018). BAL-0028 and MCC950 have no effect on NLRP1-dependent IL-18 release, but it was completely blocked by VX-765 (Fig. 2H). Together these data show that BAL-0028 specifically inhibits NLRP3 but not the AIM2, NLRC4, or NLRP1 inflammasomes.

Thus far, our data show that BAL-0028 is a specific inhibitor of NLRP3 in human myeloid cells. However, to advance BAL-0028 into *in vivo* studies in mice we needed to determine its effects in mouse cells. Surprisingly, upon activation of NLRP3 by LPS and nigericin in the mouse macrophage cell line J774A.1 the IC_50_ of BAL-0028 for IL-1β release is increased to >6 μM (Fig. 3A). This is a 114-fold increase over the IC_50_ for human THP-1s in the same assay (Fig. 1B), and contrasts with MCC950 whose IC_50_ is highly consistent between the two cell lines (5.3 nM and 14.3 nM). We hypothesized that this difference in potency of BAL-0028 could be due to species differences in NLRP3. We therefore examined the effects of BAL-0028 and MCC950 on IL-1β release in NLRP3 activation assays (LPS + nigericin) in a range of mammalian monocytes and peripheral blood mononuclear cells (PBMCs) (Fig. 3B-G). In cells from rat, dog, and rabbit BAL-0028 does not block IL-1β release, whereas MCC950 potently inhibits NLRP3 activation (Fig. 3B-D). We next examined species more closely related to humans. In cells from African Green and Cynomolgus monkeys (*Chlorocebus sabaeus* and *Macaca fascicularis*) BAL-0028 inhibits NLRP3-dependent IL-1β release with a potency similar to that observed for human myeloid cells (Fig. 3E-G compared to Fig. 1 B, E-H). BAL-0028 thus appears to be a selective inhibitor of primate NLRP3 with significantly reduced potency in other species.

**Figure 3.**
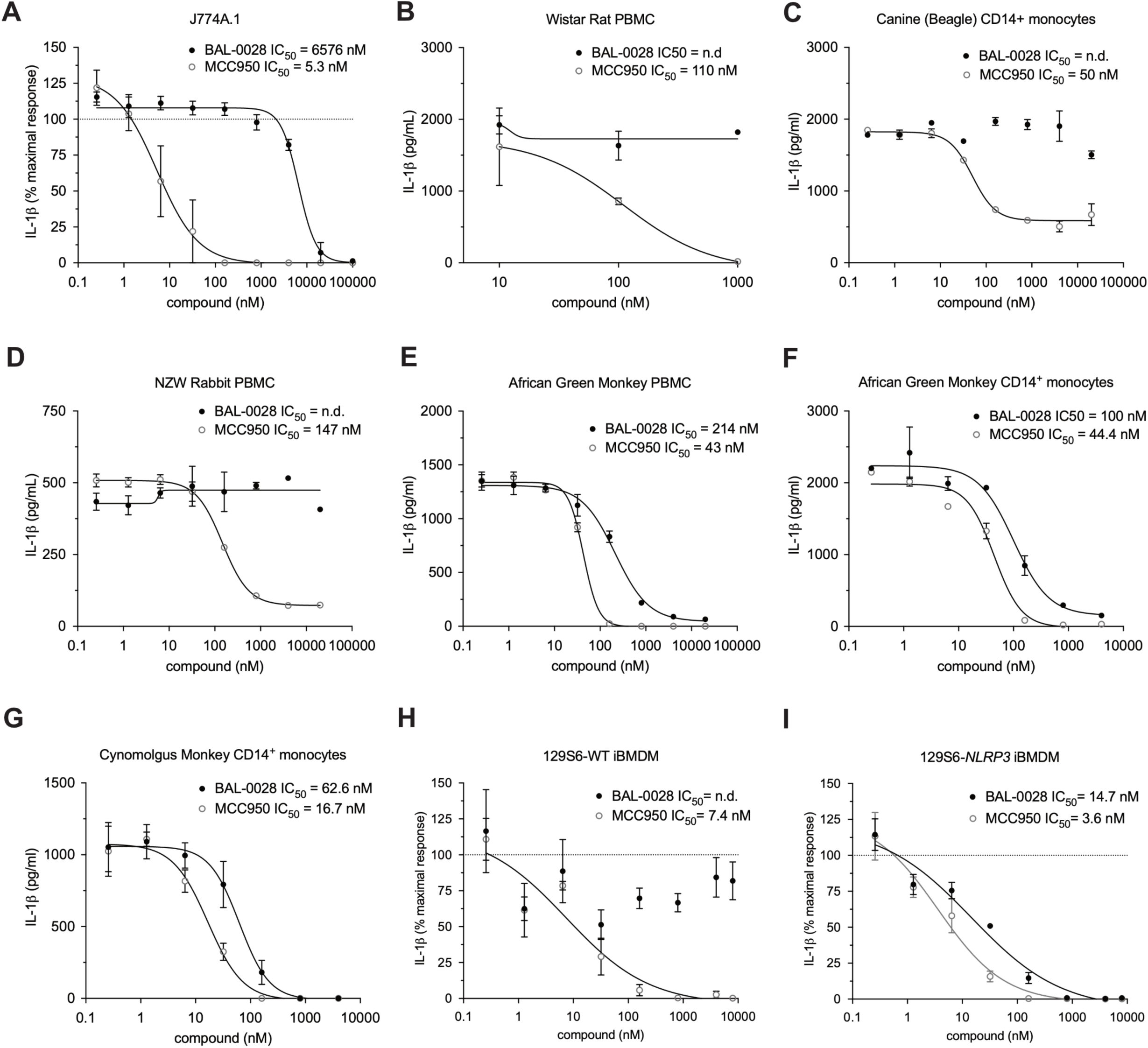
BAL-0028 inhibits primate NLRP3 but is a poor inhibitor of NLRP3 from other mammals. Comparison of BAL-0028 and MCC950 in IL-1β release assays from cells stimulated with LPS and nigericin. (A) J774A.1 mouse macrophage cell line, (B) Wistar Rat PBMCs, (C) Beagle CD14^+^ monocytes, (D) New Zealand white rabbit PBMC, African Green Monkey (*Chlorocebus sabaeus)* PBMC (E) and CD14^+^ monocytes (F), (G) Cynomolgus monkey (*Macaca fascicularis)* CD14^+^ monocytes, (H) wild-type 129S6 iBMDM and (I) 129S6-human promoter *NLRP3* iBMDM. (A, H, I) Graph symbols show average IL-1β values relative to vehicle control +/− S.E.M. from N=3 independent experiments performed in triplicate. (B-G) Graph symbols show average IL-1β values relative to vehicle control +/−S.D. from one experiment performed in duplicate (C, D,F) or triplicate (B, E, G,). IC50 curves fitted by non-linear regression analysis.

To confirm whether the difference in response to BAL-0028 observed in our human and mouse cell assays was due to species differences in NLRP3, we obtained a humanized NLRP3 mouse previously generated by Koller and colleagues where the *Nlrp3* locus is deleted and replaced with syntenic human *NLRP3* DNA including the human promoter region(Snouwaert et al., 2016). We examined BAL-0028 and MCC950 in NLRP3 assays in immortalized bone marrow-derived macrophages (BMDM) from wild-type (WT) 129S6 and 129S6-*NLRP3* mice. As expected in the WT cells BAL-0028 did not significantly inhibit IL-1β or LDH release triggered by LPS and nigericin treatment while MCC950 potently blocked NLRP3 activation (Fig. 3H, Supp. Fig 3A). However, in the *NLRP3* cells BAL-0028 inhibits NLRP3 activation, blocking both IL-1β and LDH release with an IC_50_ of 14.7 nM for IL-1β release (Fig. 3I, Supp. Fig. 3B). We observed similar results in primary peritoneal macrophages where BAL-0028 only inhibits IL-1β release in *NLRP3* cells stimulated with LPS and nigericin (Supp. Fig. 3C-D). The potency of BAL-0028 is lower than MCC950 which is consistent with our observations in human cell assays (Fig. 1B-E,I and 2B). These data indicate that the species specificity of BAL-0028 is determined by inherent differences in NLRP3 protein structure between primates and other mammals. This is a remarkable finding as mouse and human NLRP3 are highly conserved(Anderson et al., 2004; Putnam et al., 2023). To try and identify areas that may distinguish primate NLRP3 we performed a multiple sequence alignment comparing NLRP3^NACHT^ from various species (Supp. Fig. 3E). We focused on the NACHT domain since BAL-0028 was identified via a DEL screen using NLRP3 lacking the N-terminal PYD and was shown to interact with a construct lacking both the PYD and LRR domains in a surface plasmon resonance assay, indicating that BAL-0028 interacts with the NACHT domain(Hartman et al., 2024). While the amino acid sequences are generally conserved, there is a cluster of residues in the FISNA that are distinct between humans and primates relative to mouse and other mammals. Therefore, the FISNA domain may be an important determinant in how BAL-0028 interacts with NLRP3.

To examine the mechanism of action of BAL-0028 we next tested its activity in an *in vitro* ATPase assay, as many small molecule NLRP3 inhibitors such as MCC950 have been found to block NLRP3 ATPase activity(Coll et al., 2022). Consistent with previous reports(Coll et al., 2019; Lee et al., 2024; Yu et al., 2024) MCC950 inhibited the ATPase activity of recombinant NLRP3 (Fig. 4A). In contrast, BAL-0028 did not have any inhibitory effect on NLRP3 ATPase activity (Fig. 4A). This confirms that the mechanism of action of BAL-0028 is distinct from MCC950 and does not involve NLRP3 ATPase inhibition.

**Figure 4.**
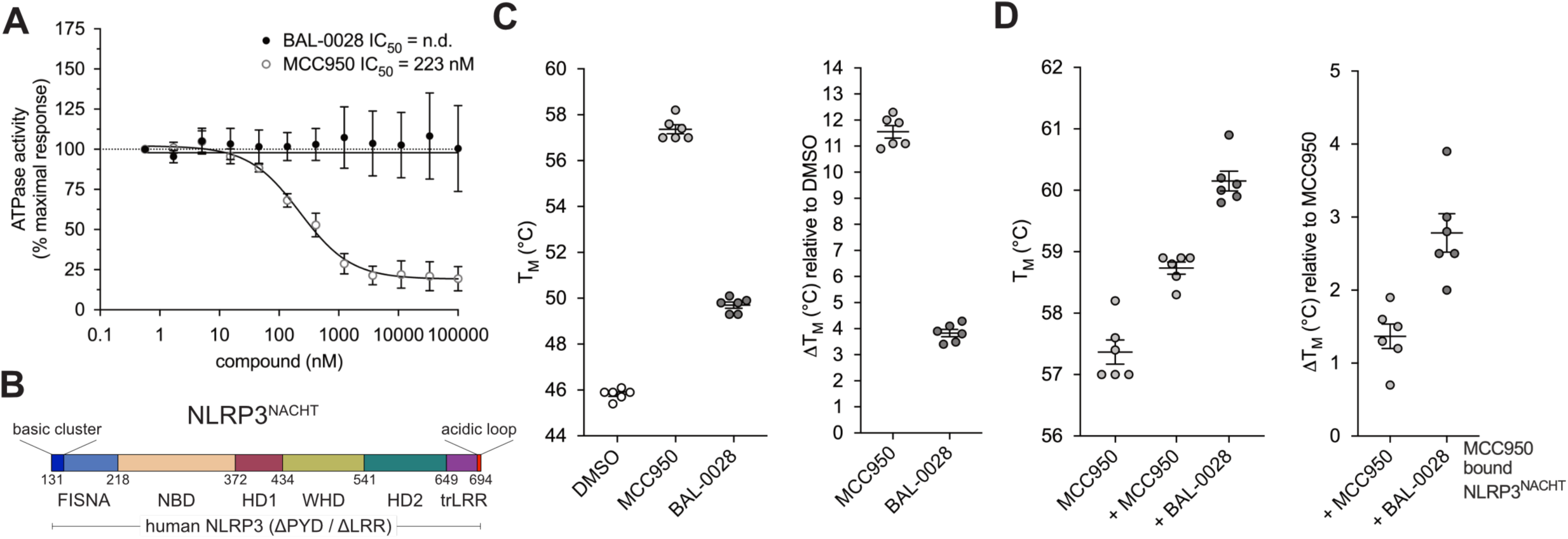
BAL-0028 does not inhibit NLRP3 ATPase activity and binds to the NLRP3 NACHT at a site distinct from MCC950. (A) Comparison of BAL-0028 and MCC950 in an ATPase activity assay with recombinant MBP-ΔNLRP3-HIS protein construct. Graph symbols show average values relative to vehicle control +/− S.E.M. from N=3 independent experiments (B) Schematic illustration of the NLRP3^NACHT^ recombinant protein used for nanoDSF studies. (C-D) NanoDSF analysis of 3 μM NLRP3^NACHT^ incubated with 10 μM BAL-0028 or MCC950 (C) or pre-incubated with 10 μM MCC950 before addition of 10 μM BAL-0028 or MCC950 (D). Graph symbols show melting temperature T_m_ or change in T_m_ relative to DMSO vehicle control or relative to MCC950 bound NLRP3^NACHT^ +/− S.E.M from N=6 independent experiments.

We next examined where BAL-0028 may be interacting with NLRP3. We performed cell-based drug affinity responsive target stability (DARTS) assays with full length human NLRP3 and NLRP3 lacking the LRR domain (NLRP3 amino acids 1-668) using BAL-0028 and MCC950. Both BAL-0028 and MCC950 prevented protease mediated degradation of NLRP3 relative to DMSO control as shown by the stabilization of bands in the immunoblots for NLRP3 (Supp Fig. 4 A-C). These data support the conclusion that BAL-0028 does not interact with the LRR and instead interacts with the central NACHT region of NLRP3.

We further investigated BAL-0028’s interaction with NLRP3 using a nano-differential scanning fluorimetry (nanoDSF) approach with recombinant NLRP3 lacking the PYD and LRR domains (NLRP3^NACHT^, Fig. 4B). Both BAL-0028 and MCC950 increase the apparent melting temperature (T_m_) of NLRP3^NACHT^ indicating that they bind to and stabilize NLRP3^NACHT^. The change in T_m_ was ∼11.5°C for MCC950 and ∼4°C for BAL-0028 (Fig. 4C). Interestingly, in a competition assay where MCC950 is prebound to NLRP3^NACHT^, the addition of BAL-0028 further stabilizes the protein with an increase in T_m_ of almost 3°C whereas additional MCC950 increases the melting temperature only by 1.3°C (Fig. 4D). These experiments demonstrate that BAL-0028 directly binds to NLRP3^NACHT^ at a site that is distinct from the MCC950 binding pocket.

BAL-0028 has very high mouse plasma protein binding with ∼99.9% of the compound bound to plasma components (Table 1) and is therefore not optimal for use in *in vivo* efficacy studies. To address this, we generated a derivative of BAL-0028, BAL-0598 (with improved pharmacokinetic properties that is approximately 16 times less mouse plasma protein bound (98.39% ± 0.49%) and therefore more compatible with *in vivo* administration (Supp. Table 1). For *in vivo* studies we elected to use 129S6 mice where human *NLRP3* is regulated by the mouse promoter, as they have increased responses to LPS compared to human promoter *NLRP3* mice (Koller et al., 2024). We first confirmed that BAL-0598 acts similarly to BAL-0028 and potently inhibits NLRP3 activation in peritoneal macrophages derived from mouse promoter *NLRP3* mice stimulated with LPS and ATP (Fig. 5A and Supp. Fig. 5A). To examine NLRP3 inhibition *in vivo* we performed a well characterized model of peritonitis using intraperitoneal (IP) injection of LPS followed by ATP where IL-1β production in the peritoneal cavity is NLRP3-dependent (Supp Fig. 5B)(Daniels et al., 2016; Pan et al., 2007; Perregaux et al., 2001). We observe a dose-dependent decrease of IL-1β in mice that received an oral dose of BAL-0598 prior to NLRP3 activation with ATP (Fig. 5B). IL-6 levels were not attenuated by any dose of BAL-0598 (Fig. 5C). The effective dose (ED_50_) of BAL-0598 to reduce the IL-1β response by half is approximately 28.6 mg/kg (Fig. 5D).

**Figure 5.**
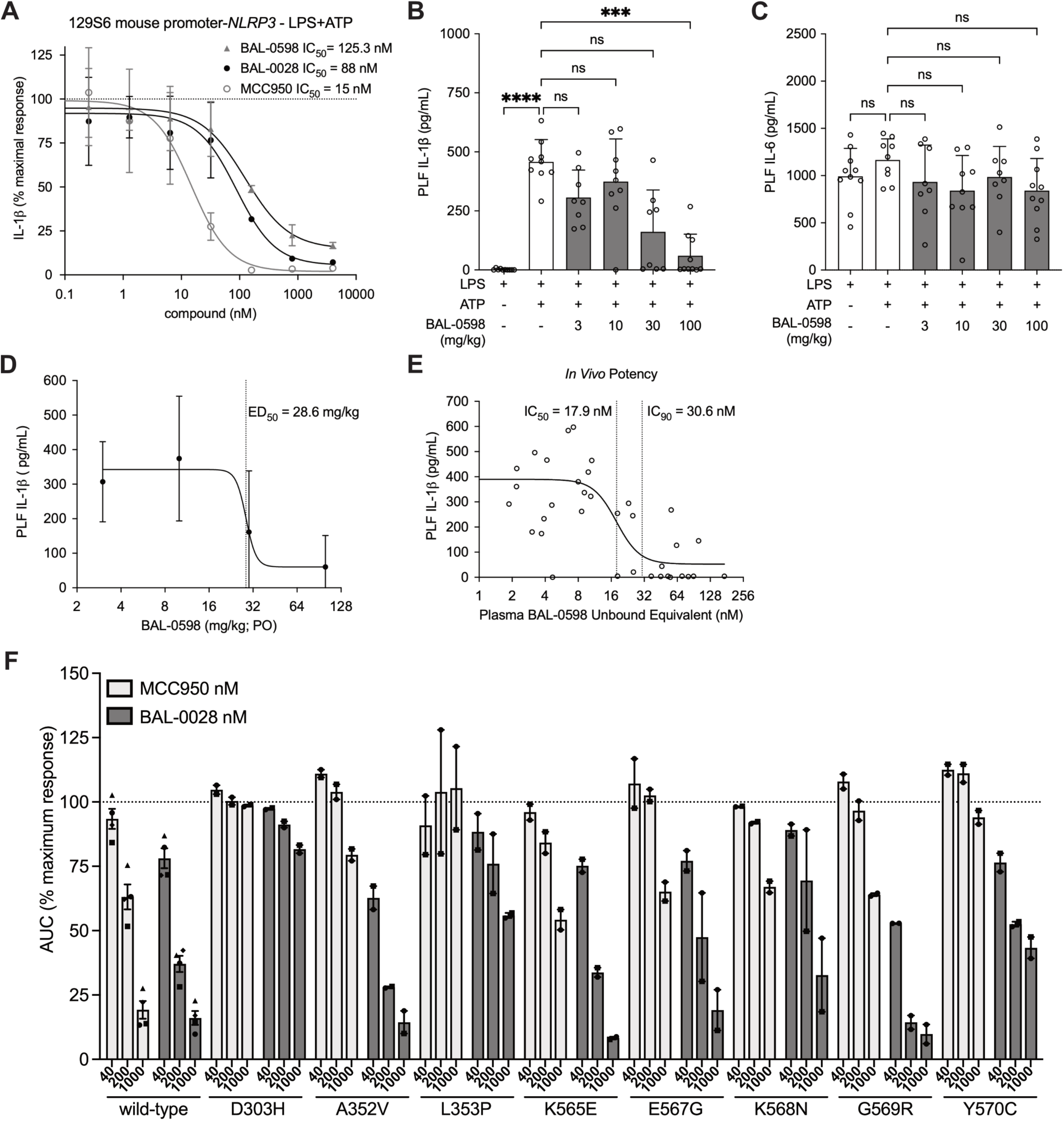
BAL-0598 inhibits NLRP3 *in vivo* and BAL-0028 inhibits NLRP3-AID mutants. (A) Comparison of BAL-0598, BAL-0028, and MCC950 in an IL-1β release assay in primary peritoneal macrophages isolated from 129S6 mouse promoter-*NLRP3* mice stimulated with LPS and ATP. Graph symbols show average IL-1β values relative to vehicle control +/− S.E.M. from N=2 independent experiments performed in duplicate. (B-C) Levels of IL-1β (B) and IL-6 (C) in the peritoneal lavage fluid of 129S6 mouse promoter-*NLRP3* mice after peritonitis induced by LPS and ATP in the presence of increasing amounts of BAL-0598. Graph symbols show values from individual mice +/− S.D. (D) Average IL-1β release values +/− S.D. in PLF from mice treated with BAL-0598 prior to peritonitis, ED50 curve fitted by non-linear regression analysis. (E) IL-1β release values in PLF from individual mice treated with BAL-0598 prior to peritonitis plotted against corresponding plasma levels of unbound BAL-0598, IC50 and IC90 curve fitted by non- linear regression analysis. (F) Comparison of BAL-0028 and MCC950 (40-1000 nM) in a cell death assay in U937 cells expressing NLRP3 or NLRP3-AID mutants (D303H, A352V, L353P, K565E, E567G, K568N, G569R, Y570C) stimulated with LPS and nigericin. Graph symbols show average area under the curve (AUC) values relative to DMSO vehicle control for each cell type +/− S.E.M. from N=2-4 independent experiments performed in duplicate.

**Table 1.** Mouse plasma protein binding of BAL-0028 and BAL-0598.

To better understand how systemic exposure of BAL-0598 correlates to IL-1β release in the peritoneal cavity, *in vivo* potency was determined by plotting the unbound plasma levels of BAL-0598 against the absolute IL-1β concentrations measured in the peritoneal lavage fluid (PLF) of individual mice. Just prior to lavage of the peritoneal cavity, terminal plasma samples were collected, and the total plasma concentrations of BAL-0598 were determined by LC-MS/MS (Supp Fig. 5C). The unbound BAL-0598 equivalent for each mouse was then calculated using the mean unbound protein plasma binding value of 1.61% (Supp. Table 1). The *in vivo* dose-response inhibition curve indicates that after oral dosing, the unbound systemic exposure of BAL-0598 required to reduce the IL-1β response by half is approximately 17.9 nM, and by 90% is 30.6 nM (Fig. 5E). Importantly, the *in vivo* potency data indicates that the ability of BAL-0598 to inhibit peritoneal macrophages activated with LPS and ATP *ex vivo* underestimates the inhibitory activity of BAL-0598 *in vivo* (Fig. 5A).

Finally, to examine whether BAL-0028 or its derivatives could be used to treat disease-associated NLRP3 mutations we employed a U937 cell model to express gain-of-function NLRP3-AID mutants as previously described(Cosson et al., 2024) (Supp Fig. 5E). NLRP3 or NLRP3-AID mutant expression is induced by doxycycline treatment in U937s and NLRP3 is activated by stimulation with LPS and nigericin. We selected eight NLRP3-AID gain-of-function mutations including known mutations shown to be relatively insensitive to MCC950 inhibition (D303H, L353P, G569R) and well characterized constitutively active mutants (A352, K565E, E567G, K568N, Y570C)(Cosson et al., 2024). Pre-treatment of wild-type NLRP3 with MCC950 or BAL-0028 (40-1000 nM) dose-dependently inhibits cell death (Fig. 5F). However, in NLRP3 mutant cells MCC950 fails to inhibit (D303H, L353P) or partially inhibits cell death at 1000 nM (A352V, K565E, E567G, K568N, G569R, Y570C), as expected. In contrast, BAL-0028 potently inhibits A352V, K565E, E567G, K568N, G569R at levels similar to wild-type NLRP3, and partially inhibits D303H, L353P, and Y570C at 1000 nM. These data demonstrate the ability of BAL-0028 to inhibit disease associated NLRP3 mutations, and particularly those that may be insensitive to MCC950-like small molecules.

## DISCUSSION

Previous efforts to identify NLRP3 inhibitors have largely focused on cell based phenotypic screens in mouse and human myeloid cells. As NLRP3 activation is a highly complex process involving multiple priming and activation mechanisms for different stimuli(McKee and Coll, 2020; Swanson et al., 2019), deconvolving ‘hits’ from such cellular screens can be extremely challenging. We therefore chose an alternative approach and employed a DEL screen to identify compounds that interact with recombinant NLRP3, yielding the lead compound BAL-0028(Hartman et al., 2024). Here we have characterized the mode of action of BAL-0028 in depth, benchmarking it with the tool molecule MCC950. We observe that BAL-0028 inhibits NLRP3 signaling in a range of human cell types and in response to multiple NLRP3 stimuli (Figs. 1-2). BAL-0028 is a specific inhibitor of NLRP3 as it does not prevent activation of the NLRP1, NLRC4 or AIM2 inflammasomes and does not affect LPS-induced signaling (Supp Fig 1, Fig 2). In these human cell-based NLRP3 activation assays, BAL-0028 effectively phenocopies MCC950 albeit at slightly higher concentrations.

Surprisingly, when we examined BAL-0028 in mouse macrophages we noticed a highly significant decrease in its ability to inhibit NLRP3 (Fig. 3). A recent study also described an NLRP3 inhibitor NDT-0796 that is active in human cells but inactive in mouse macrophages(Smolak et al., 2024). This difference was because NDT-0796 is a pro-drug that must be metabolized by carboxylesterase-1 (CES-1) into its active form, and mouse macrophages do not express CES-1(Smolak et al., 2024). As BAL-0028 does not contain a carboxylate this explanation for the species differences observed was unlikely. A molecule termed J114 that inhibits both NLRP3 and AIM2 inflammasomes is also reported to be significantly less potent in mouse macrophages relative to human cells(Jiao et al., 2022). However, the reason for this species difference was not examined. We chose to explore the ability of BAL-0028 to inhibit NLRP3 in other species, and while BAL-0028 could not inhibit rat, dog, or rabbit NLRP3, it potently blocked monkey NLRP3 (Fig. 3). BAL-0028 is thus active in humans and closely related monkey species, making it the first primate specific NLRP3 inhibitor to be reported. This species specificity is surprising given the high sequence conservation of the NACHT domain in mammalian NLRP3 (Anderson et al., 2004), but is consistent with the use of human NLRP3 protein in the DEL-screening assay. In line with this, our studies using 129S6 mice expressing human NLRP3 unequivocally show that the potency of BAL-0028 is dependent on the differences between mouse and human NLRP3 (Figs. 3 and 5). Thus, even subtle changes in NLRP3 structure between mouse and human can be exploited for drug development. While much of our knowledge of inflammasome biology has been informed by mouse models there are many differences in inflammasome structure, expression, and regulation in humans. For example, NLRP1 biology has significantly advance in recent years due to studies on keratinocytes which express NLRP1 in humans but not in mice. NLRP1 is also structurally distinct in mouse where it lacks a PYD domain(Barry et al., 2023). There are also two families of regulatory proteins termed Pyrin only proteins (POPs) and CARD-only proteins (COPs) that only arose in the primate lineage. COPs and POPs are known to regulate inflammasome formation and signaling, but functional studies have been relatively limited as they are not expressed in mice(Devi et al., 2020). Our work highlights the importance of performing screens using human proteins and cell systems to capture these important biological differences in the immune system.

The species-specific effects of BAL-0028 also reveal a clear difference with MCC950, which inhibited NLRP3 in all species tested. Our mechanistic studies further show that BAL-0028 does not inhibit the ATPase activity of NLRP3 distinguishing it from MCC950 and many other NLRP3 inhibitors such as CY-09 that appear to converge on inhibiting NLRP3 via blocking this ATPase activity(Coll et al., 2022). We conclusively show that BAL-0028 binds to NLRP3 in the NACHT domain using DARTS and nanoDSF assays. Indeed, our nanoDSF analysis reveal a novel synergistic stabilization of NLRP3^NACHT^ by BAL-0028 in the presence of MCC950 showing that they bind at different sites of the protein. BAL-0028 therefore inhibits NLRP3 by a mechanism distinct from MCC950 and many other previously characterized NLRP3 inhibitors. Based on our analysis of NLRP3 sequences we speculate that BAL-0028 may interact in the FISNA subdomain of the NLRP3^NACHT^, although it is not currently understood how this proposed binding interaction is consistent with the computer-aided drug design (CADD) binding site prediction previously published for BAL-0028(Hartman et al., 2024). While we have not experimentally identified the specific binding site of BAL-0028, this is the subject of ongoing structural biology studies.

For the future development of BAL-0028 related compounds establishing *in vivo* efficacy is important. As a first step we demonstrated that BAL-0028 is active in *ex vivo* human whole blood NLRP3 assays (Fig. 1); however, we developed an analogue BAL-0598 for *in vivo* studies due to the high plasma protein binding of BAL-0028. In the human NLRP3 129S6 mice in a peritonitis model we show that BAL-0598 dose dependently blocks IL-1β release but not IL-6 (Fig. 5). Furthermore, our *in vivo* peritonitis studies indicate that *ex vivo* cell-based potency assays performed in peritoneal macrophages underestimated the *in vivo* potency after oral dosing of BAL-0598, based on the *in vivo* dose-response inhibition curve derived from unbound BAL-0598 plasma values and absolute IL-1β in the PLF.

Using a cell-based model of NLRP3-AID we show that BAL-0028 can also inhibit gain-of-function NLRP3 mutations. All the mutations we examined are in the NACHT within the nucleotide binding domain (NBD) or helical domain (HD2) causing constitutively active or hyperactive NLRP3 variants. A number of these mutations (D303H, K568N, Y570C) are known to cause the severe NLRP3-AID neonatal-onset multisystem inflammatory disease (NOMID) (Infevers Database) and include several somatic mutations (D303H, K565E, E567G, K568N, Y570C)(Cosson et al., 2024). BAL-0028 was able to fully or partially inhibit these highly pathogenic NLRP3 mutations, suggesting that BAL-0028 or its derivatives can be effective treatments for many NLRP3-AID disease variants. Importantly, for all these mutations BAL-0028 was a more effective inhibitor than MCC950. BAL-0028 or its derivatives may thus be an alternative therapeutic modality for NLRP3-AID patients that may not respond to MCC950-derived molecules.

In summary we have characterized a novel small molecule inhibitor that is potent and specific for primate NLRP3. We anticipate that future work on BAL-0028 will reveal novel insights into the biology of NLRP3 and that improved understanding of the structural biology of human NLRP3 will inform the development of BAL-0028 related compounds.

## MATERIALS AND METHODS

### Source of BAL-0028 and MCC950

The synthesis of BAL-0028 is described in detail in Hartman et al.(Hartman et al., 2024). MCC950 was purchased from Adipogen, ApexBio (Houston, TX) or Selleck Chemicals (Houston, TX).

### THP-1 Cell-Based Assays

Human monocytic THP-1 cells (ATCC) were cultured in growth media (THP GM) until they reached logarithmic growth and achieve a viability > 90%. THP GM is composed of RPMI-1640 + GlutaMAX (Gibco)/ 10% fetal bovine serum (Corning)/ 55 µM β-mercaptoethanol (Gibco)/ pen/strep (Caisson). Cells were spun down and resuspended to 1,000,000 cells/mL in THP GM containing either 20 nM or 500 nM PMA (Sigma). 150,000 cells (150 µL) were then added to each well of a 96-well TC plate and incubated for either 2 4hr or 3 hr, respectively, in a standard cell culture incubator (37°C; 5% CO_2_). In the NLRP3 inflammasome activation assays, 200 µL of THP GM or THP GM containing 100 ng/mL LPS (*E.coli* O26:B6; Sigma) was then added to the appropriate wells and the cells incubated an additional 3 hrs. The media was then replaced with Opti-Mem medium (without serum; Thermo) containing pre-determined dilutions of test compounds in replicate wells. After a 30 min incubation, 10 µM nigericin (final concentration; Sigma) in Opti-Mem medium with the corresponding concentration of test compound was added to the wells for an additional 1 hr. Positive control wells contain 10 µM nigericin in Opti-Mem in the absence of test compound, while negative control wells contain Opti-Mem only. Supernatants were then assayed for IL-1β (human; DuoSet; R&D) and relative LDH levels (as a surrogate for pyroptosis) using a CytoTox 96 Kit (Promega). For the THP-1 LPS signaling assays, 200 µL of THP GM (negative control wells), THP GM containing 100 ng/mL LPS (positive control wells) or THP GM containing 100 ng/mL LPS with pre-determined dilutions of test compounds in replicate wells was added to the appropriate wells for 3 hr. Supernatants were then assayed for IL-6 (human; DuoSet; R&D), TNF-α (human; DuoSet; R&D) and relative LDH levels using a CytoTox 96 Kit (Promega). For THP-1 viability assays, after the PMA incubation step, the media was replaced with Opti-Mem medium (without serum; Thermo) or Opti-Mem medium containing pre-determined dilutions of test compounds in replicate wells and incubated for 1.5 hr. Supernatants were then assayed for relative LDH levels using a CytoTox 96 Kit (Promega). In all THP-1 assays, once supernatants were removed, the relative viability of adherent cells in the 96-well TC plate were determined using a CellTiter-Glo luminescent cell viability assay (Promega).

### Human Microglial NLRP3 inflammasome activation assay

Human induced-pluripotent stem cell-derived microglia (iCell Microglia; R1131, Fujifilm) were directly thawed in poly-L-lysine coated 96-well plates at ∼50,000 cells/well in iCell glial base medium supplemented with iCell microglia supplements A, B, and C as per manufacturer’s instructions. Cells were incubated at 37 °C, 5% CO_2_. Next day, 50% of the glial medium was replaced with fresh medium and cells were incubated for a further 2 days. For potency determination, iCell Microglia were first primed with 200 ng/mL LPS for 4 hours and then pre-incubated for 30 min with 5-fold compound concentrations ranging from 0.256 nM to 4 µM. Cells were then stimulated with 10 µM nigericin with the corresponding compound concentrations and incubated for an additional 30 minutes. Supernatants were collected for cytotoxicity assay and cytokine ELISAs. Cells treated with 200 ng/mL LPS and 10µM nigericin (positive control) and cells with medium alone (negative control) were included in the experiment. MCC950 was used as a tool compound.

### Human Monocyte-derived Macrophages (HMDM) NLRP3 inflammasome activation assay

Buffy coats were obtained from the Northern Ireland Blood Transfusion Service (NIBTS reference number 2019/05), ethical approval for the use of buffy coats was from the Queen’s University Faculty Research Ethics Committee (reference MHLS 19_17). Peripheral blood mononuclear cells were isolated by density centrifugation using Ficoll-Paque Plus (GE Healthcare, 12440053). CD14+ monocytes were then isolated using magnetic-activated cell sorting CD14+ positive selection (Miltenyi Biotech, 130-050-201) according to the manufacturer’s instructions. CD14+ monocytes were cultured in RPMI 1640, 10% FCS, 50 U/ml pen/strep (all Gibco) and 50 ng/ml recombinant human macrophage colony-stimulating factor (M-CSF) (ImmunoTools, 11343118) for seven days. HMDM were seeded at 0.7 ×10^6^ cells/ml in 96-well TC plates.

For NLRP3 assays, the following day the media was removed and replaced with OptiMem +/− 100 ng/mL Ultrapure LPS from Escherichia coli K12 (Invivogen) for 4 hr. Inhibitors were prepared by serial dilution in DMSO. The media were removed and replaced with OptiMem containing inhibitors or 0.1% DMSO vehicle control. After 30 min incubation 5 μM Nigericin (Adipogen) was added for 2 hr. Cell free supernatants were removed and IL-1β and TNF were assessed by ELISA according to the manufacturer’s instructions (Thermo Fisher, 88-7261-77, 88-7346-77).

### Human induced pluripotent stem cell (iPSC)-derived macrophages (iMacs) NLRP3 inflammasome activation assay

KOLF2-C1 iPSCs were obtained from the Wellcome Sanger Institute (Hinxton, UK). iPSCs were cultured on vitronectin-coated (ThermoFisher, A14700) plates in Essential 8 medium (ThermoFisher, A1517001) at 37°C in a humidified 5% CO2 incubator with 10 μM Rho-kinase (Rock) inhibitor Y-27632 Dihydrochloride (Merck, Y0503) added following thawing and passaging. Embryoid bodies (EBs) were formed as previously described(Douthwaite et al., 2022). In brief, iPSCs were seeded at 1 × 10^5^ cells in 100 μL per well onto low-adherent, U-bottom 96 well plates in Essential 8 medium supplemented with 50 ng/mL BMP-4 (R&D Systems, 314-BP010) and 10 μM Rock inhibitor. The plates were centrifuged for 3 min at 800 × g, 4°C and incubated for 5 days. The EBs were then transferred to gelatine-coated (Sigma-Aldrich, G1890) tissue culture flasks in myeloid precursor base medium: X-VIVO 15 Serum Free Medium (Lonza, BE02-060F), 1 × GlutaMAX and 1% P/S. Media was changed every 4–5 days and the EBs began to produce myeloid precursor cells approximately 3–4 weeks after transferring to the gelatine-coated flasks.

To harvest the precursor cells, the medium was collected from the EB flasks and centrifuged for 3 min at 300 × g and the cells resuspended in iMac medium: RMPI 1640, 10 % FBS, 1 × GlutaMAX and 1% Pen/Strep supplemented with 100 ng/mL M-CSF (Proteintech, HZ-1192). Myeloid precursor cells were differentiated for 7 days then harvested using Lidocaine monohydrate (Sigma-Aldrich, L5647) and plated onto 96-well TC plates at 0.5 × 10^6^ /mL in 100 μL per well in iMac medium.

NLRP3 assays were conducted as described for HMDM above except cytotoxicity was also assessed in cell free supernatants by LDH assay (Takara Bio, MK401).

### Human Whole Blood Assays

The human whole assays followed approximately the procedure described previously (Grinstein et al., 2018). Human blood sample was collected by a licensed phlebotomist in sodium heparin-containing tubes (WCG IRB Protocol #20223689). 140 µL of heparinized whole blood was then added to each well of a 96-well TC plate. 20 µL of PBS (no Ca^2+^ and no Mg^2+^), PBS containing 8 µg/mL LPS (*E.coli* O26:B6; Sigma; final concentration 1 µg/mL) or PBS containing 8 µg/mL LPS (final concentration 1 µg/mL) with pre-determined dilutions of test compounds in replicate wells was added to the wells for 3 hr on a shaker (180 rpm) in a standard cell culture incubator (37°C; 5% CO_2_). After 3 hr, 40 µL of 5 mM ATP (Sigma-Aldrich, A6419; final concentration 1 mM) was added to the appropriate wells and incubated for an additional 30 min on a shaker (180 rpm). At the end of all treatments, 100 μl of PBS was added to each well, the plates centrifuged at 450 xg for 15 min at RT and the supernatant from each well was assayed for IL-1β (human; DuoSet; R&D) or IL-18 (human; DuoSet; R&D).

### Western Blots

THP-1 cells are cultured as described above with the following differences: 4.5 x 10^6^ cells were seeded in THP GM containing 500 nM PMA (Sigma) into each well of a 6-well cell culture plate and incubated for 3hr. After this incubation, the plate was tilted and media carefully removed. 2mL of THP GM or THP GM containing 100 ng/mL LPS (*E.coli* O26:B6; Sigma) is then added to the appropriate wells and the cells incubated an additional 3 hrs. The media is again removed and replaced with Opti-Mem medium (without serum; Thermo) OR Opti-Mem medium (without serum; Thermo) containing 200 - 500 nM of test compound. After a 30 min incubation, 10µM nigericin (final concentration; Sigma) in Opti-Mem medium with 200 - 500 nM of test compound is added to the wells for an additional 30 min. Positive control wells contain 10µM nigericin in Opti-Mem in the absence of test compound, while negative control wells (PMA or PMA + LPS treated cells) contain Opti-Mem only. Supernatants are collected and condensed 10-fold by centrifugation using Pierce Protein Concentrator columns (Thermo; 88512) according to the manufacturers supplied instructions. NuPAGE LDS sample buffer (Thermo; NP0007) with reducing agent (Thermo; NP0009) are then added to the appropriate volume of condensed supernatant samples for gel loading. The adherent cells were then washed 2 x with ice cold TBS, all liquid was subsequently removed and the cells lysed with 100 µL per well of 1x RIPA buffer (BioBasic RB4475) containing protease cocktail (Sigma; P8340) and caspase-1/4 inhibitor VX-765 (Invivogen, inh-vx765i-1) while on ice. Insoluble cellular material was removed by cold centrifugation at 16,000xg. The lysates were then resuspended with 1x RIPA buffer and sample buffer with reducing agent to a final concentration of 0.4 mg/mL. 10 µg of protein as loaded per well of a 10% Tris-Bis gel (Thermo; NP0301BOX) and run at 100V for approximately 2-3 hours, after which the proteins were transferred to PDVF membrane using a Iblot (IB1001) transfer apparatus at 20V for 7 min. Membranes were then blocked in TBS-T (10 mM Tris-HCl (pH 7.4), 150 mM NaCl, 0.2% Tween-20) containing 5% BSA (VWR; AAJ65097-22) at RT for 45 min while shaking. Membranes were then incubated with one of the following primary antibodies: NLRP3 (Cell Signaling; 15101), cleaved IL-1β (Cell Signaling; 12242), cleaved CASP1 (Cell Signaling; 4199) and actin (Sigma; A2066). After incubation, the membranes were washed and incubated with one of the compatible secondary antibodies: anti-mouse (Jackson, 115-035-062) or anti-rabbit (Jackson, 111-035-144) in TBS-T. Proteins were detected by incubating the membrane with Femto SuperSignal substrate (Thermo; 34095) and luminescence signal was detected using a Bio-Rad ChemiDoc XRS+ Imaging System.

### ASC Speck Assays

THP-1-ASC-GFP (InvivoGen) were maintained in growth media containing RPMI-1640 + GlutaMAX (Gibco) with 10% FBS (Corning) and 1% penstrep (Caisson) at 37°C in 5% CO2 atmosphere. Cells were seeded in a 96-well plate (150,000 cells/well) in growth media containing 100 nM PMA for 3 hr. PMA-treated cells were primed with 100 ng/mL LPS (O6:B26; Sigma-Aldrich) for further 3 hr, followed by a 30 min preincubation with BAL-0028 or MCC950, at 5-fold concentrations ranging from 0.256 nM to 20 µM. To activate NLRP3, cells were stimulated with 10 µM nigericin with and without inhibitors for 30 min. Nuclei were stained with the live stain, Hoechst 33342 for 5 min. Cells were imaged live (20X magnification) and analyzed using the CellInsight CX5 high content screening platform (Thermo Fisher Scientific). Each well was imaged in two channels (GFP and DAPI) for 9 fields per well. The number of nuclei (DAPI) and perinuclear ASC specks (GFP) were counted and percent ASC specks/nuclei were calculated. For IC_50_ determination, the % ASC speck positive cells were plotted against BAL-0028 or MCC950 concentrations (nM).

### HEK293T cell ASC speck formation assay

HEK293T cells stably expressing ASC-BFP fusion protein were used as previously described (Hochheiser et al., 2022a). Cells were seeded into 24-well TC plates at a density of 125,000 cells per well and incubated overnight at 37°C. To induce moderate expression of NLRP3 for assessable ASC speck formation, cells were transfected with 100 ng per well of a doxycycline-inducible TetO6-NLRP3-hPGK-TetON3G-T2A-mCherry construct using Lipofectamine2000 (ThermoFisher Scientific) according to the manufacturer’s instructions. Sixteen hours post-transfection, NLRP3 expression was induced by adding doxycycline (10 ng/ml). Simultaneously, cells were treated with increasing concentrations of BAL-0028 or MCC950 and incubated for 4 hours. To induce ASC speck formation, cells were subsequently treated with 10 µM nigericin for 1 hr.

After trypsinization cells were washed and resuspended in flow buffer (DPBS, 2 mM EDTA, 0.5% BSA). Flow cytometry was conducted at a LSRFortessa II. Gates were set to select for single cells expressing ASC-BFP. From these, mCherry-positive, NLRP3 expressing cells were selected. Among the mCherry-positive cells, the proportion of ASC speck formation was calculated as previously described by Sester et al.(Sester et al., 2015). The difference in ASC speck formation between nigericin-stimulated and unstimulated cells was plotted against increasing antagonist concentrations. These values were normalized to the speck formation difference calculated for cells without compound treatment (% maximum response). The generated dose-response curve was fitted using the four-parameter logistic equation built into Prism 10.1.1. to determine the half maximal inhibitory concentrations of BAL-0028 and MCC950.

### iMacs ASC Speck Staining and Imaging

Cells were plated at 1×10^5^ per well overnight in 24 well plates with coverslips in 500 μL iMac medium. LPS (100 ng/mL) was used to prime the cells for 3.5 h and MCC950 (1 μM) or BAL-0028 (1 μM, 100 nM, or 10 nM) were added for 30 minutes prior to the addition of nigericin (5 μM) for 2 h. Cells were washed in PBS, fixed in 4% paraformaldehyde (PFA) for 15 minutes, and stored in PBS until staining. Coverslips were washed 3 times in PBS then permeabilised with 0.1% Triton X-100 in PBS for 10 min. Coverslips were washed 3 times in PBS then 50 mM ammonium chloride was added for 10 min to quench the PFA. Coverslips were washed 3 times in PBS then blocked with 0.5% BSA/PBS for 30 min. Primary anti-ASC antibody (BioLegend, 676502) was diluted 1 in 50 in 0.5% BSA/PBS and added for 60 minutes then coverslips were washed in PBS 4 times. Alexa Fluor™ 488 Donkey anti-mouse (Invitrogen, A21202) was diluted 1 in 800 in 0.5% BSA/PBS and added for 1 hr. The coverslips were washed 4 times in PBS before mounting in Vectashield Antifade Mounting Medium with DAPI (2BScientific, H-1299) on glass microscope slides. The mounted coverslips were sealed with clear nail polish and stored at 4°C until imaging. Coverslips were imaged using the Leica LAS X software and the DM5500 fluorescent microscope at 20X and 40X. Six fields of view were analysed per condition in each experiment with 2-3 technical replicates (coverslips) for each biological replicate. ASC specks were quantified using Fiji. An automated system was used to quantify the number of DAPI+ cells and the ASC specks were manually counted. The percentage of DAPI+ASC+ positive cells was calculated.

### AIM2, NAIP/NLRC4 and NLRP1 Inflammasome Activation Assays

Monocytic THP-1 cells were seeded in 96-well plates (150,000 cells/well) in growth media containing 100 nM PMA for 3 hr. For NAIP/NLRC4, growth media was aspirated and 180 µL OptiMEM with 10-fold BAL-0028 or MCC950 (Selleck Chemicals) concentrations ranging from 10 nM to 10 µM was added to the cells. Then, 20 µL of 10X ‘NeedleTox’ comprising of a mixture of 5 µg/mL LfN-Needle (Invivogen) and 5 µg/mL anthrax toxin’s protective antigen (PA; EMD Millipore) was added to activate NAIP/NLRC4 (final concentration 5 ng/mL). Cells treated with NeedleTox (positive control) and cells with OptiMEM alone (negative control) were included in the experiment. For AIM2, PMA-treated THP-1 cells were primed with 100 ng/mL LPS (O6:B26; Sigma-Aldrich) for 24 hr. Cells were then incubated with the AIM2 ligand, PolydA:dT/Lyovec ( Invivogen; final concentration 5 µg/mL) with and without BAL-0028 or MCC950, 10-fold concentrations ranging from 10 nM to 10 µM. For NLRC4 and AIM2 inflammasomes, cells were incubated at 37°C, 5% CO_2_ for 24 hr and supernatants were collected for cytokine analysis. The caspase-1 inhibitor, VX-765 (Invivogen; final concentration 10 µM), was used as a positive control for inflammasome inhibition.

For NLRP1, human keratinocytes (NHEK; Lonza) were used for testing potency of BAL-0028. Cells were plated at a density of 40,000 cells/well in KBM™ Gold Keratinocyte Growth Basal Medium (Lonza) supplemented with KGM™ Gold Keratinocyte Growth Medium SingleQuots™ Supplements and Growth Factors (Lonza) and 0.03 µM calcium carbonate (CaCl_2_; Sigma-Aldrich). After overnight incubation at 37°C, 5% CO_2_, differentiation of NHEKs was induced by increasing the concentration of CaCl_2_ to 1.5 mM and cells were incubated for a further 24 hr. Talabostat (MedChem Express; final concentration 10 µM) was used to activate NLRP1 and cells were incubated overnight with the agonist in the presence or absence of BAL-0028 or MCC950, 10-fold concentrations ranging from 10 nM to 10 µM. Supernatants were collected for cytokine analysis. The caspase-1 inhibitor, VX-765 VX-765 (Invivogen; final concentration 10 µM), was used as a positive control for inflammasome inhibition.

### Mouse J774A.1 NLRP3 inflammasome activation assay

J774A.1 cells (ATCC) were cultured in complete growth media composed of DMEM high glucose (Gibco)/ 10% fetal bovine serum (Corning)/ pen/strep (Caisson). Cells were spun down and resuspended to 500,000 cells/mL in GM-2. 100,000 cells (200 µL) were then added to each well of a 96-well TC plate and incubated O/N in a standard cell culture incubator (37°C; 5% CO_2_). After this incubation, the plate was tilted and media carefully removed. 200 µL of GM-2 or GM-2 containing 100 ng/mL LPS (*E.coli* O26:B6; Sigma) was then added to the appropriate wells and the cells incubated an additional 5 hrs. The media was again removed and replaced with Opti-Mem medium (without serum; Thermo) containing pre-determined dilutions of test compounds in replicate wells. After a 30 min incubation, 10 µM nigericin (Sigma) in Opti-Mem medium with the corresponding concentration of test compound was added to the wells for 1 hr. Positive control wells contain 10 µM nigericin in Opti-Mem in the absence of test compound, while negative control wells contain Opti-Mem only. Supernatants were then transferred to a 96-well plate for storage and assayed for IL-1β (human; DuoSet; R&D) and relative LDH levels using a CytoTox 96 Kit (Promega). Once supernatants were removed, the relative viability of adherent cells in the 96-well TC plate were determined using a CellTiter-Glo luminescent cell viability assay (Promega).

### CD14+ (Human, Canine and Monkey) and PBMC (Minipig, Rabbit and Rat) NLRP3 inflammasome activation assays

Human CD14+ monocytes were freshy isolated from approximately 30mL of heparinized whole blood collected by venipuncture from healthy human volunteers was collected into heparin tubes from each donor. PBMCs were isolated from whole blood using Sepmate (STEMCELL Technologies; Cat No. 85450) tubes, following the manufacturer’s recommended protocol. PBMCs were then magnetically sorted to enrich CD14+ cells using microbeads conjugated to a monoclonal human CD14 antibody (Miltenyi Biotec; Cat No. 130-050-201). Cryopreserved Beagle canine CD14+, Cynomolgus monkey CD14+ cells, New Zealand white rabbit PBMCs and Wistar rat PBMCs were obtained from IQ Biosciences. Cryopreserved Gottingen minipig PBMCs were obtained from BioIVT. African green monkey (AGM-1) CD14+ cells were isolated from cryopreserved PBMCs (Virscio) using non-human primate CD14+ MACS® MicroBeads and the supplied MACS® separation protocol (Miltenyi). AGM-1 CD14+ cells were isolated immediately upon thaw of the PBMCs. Upon thaw, cells were spun down and resuspended to 500,000-700,000 cells/mL in pre-warmed complete growth composed of RPMI-1640 + GlutaMAX (Gibco)/ 10% fetal bovine serum (Corning)/ pen/strep (Caisson). 50,000-70,000 cells (100µL) were then added to the wells of a 96-well TC plate and the cells incubated 1 hr in a standard cell culture incubator (37°C; 5% CO_2_) to recover. 100 µL of GM-3 containing 200 ng/mL LPS (*E.coli* O26:B6; Sigma) was then added to the wells (100 ng/mL final concentration) and the cells incubated 5 hrs. The media was again removed and replaced with Opti-Mem medium (without serum; Thermo) containing pre-determined dilutions of test compounds in replicate wells. After a 30 min incubation, 10µM nigericin (final concentration; Sigma) in Opti-Mem medium with the corresponding concentration of test compound was added to the wells for an additional 1 hr. Positive control wells contain 10µM nigericin in Opti-Mem in the absence of test compound, while negative control wells contain Opti-Mem only. Supernatants are then transferred to a fresh 96-well plate for storage and assayed for IL-1β (canine, monkey, porcine, rabbit or rat; DuoSet; R&D) and relative LDH levels using a CytoTox 96 Kit (Promega). Once supernatants are removed, the relative viability of adherent cells in the 96-well TC plate are determined using a CellTiter-Glo luminescent cell viability assay (Promega).

### Immortalized bone-marrow derived macrophages (BMDM) NLRP3 inflammasome activation assays

Generation of 129S6 human promoter-*NLRP3* mice has been previously described(Snouwaert et al., 2016). In brief, tibias and femurs from wild-type 129S6 or 129S6 human promoter-*NLRP3* male mice were removed, and the bone marrow was harvested by flushing with fresh medium. Bone marrow was plated in BMDM medium: DMEM high glucose (Gibco), 10% FBS (Gibco), 100 mM Sodium Pyruvate (Gibco), 50 U Pen/Strep (Gibco), supplemented with 100 ng/mL human M-CSF (Proteintech). Cells were incubated in a standard cell culture incubator (37°C; 5% CO_2_) for 5 days. Media were removed and replaced with media containing J2 CRE virus (carrying v-myc and v-Raf/v-Mil oncogenes, kindly donated by Dr Joana Sá-Pessoa, Queen’s University Belfast) for 24 hr before being replaced with fresh BMDM medium containing M-CSF. Cells were continuously cultured for 9 weeks with a gradual reduction in M-CSF until cells doubled every 24 hrs in M-CSF free media. Immortalized BMDM (iBMDM) were plated at 0.5 X 10^6^ /mL in 96 well TC plates in 100 μl BMDM medium (no M-CSF).

For NLRP3 assays, the following day the media was removed and replaced with OptiMem +/− 100 ng/mL Ultrapure LPS from Escherichia coli K12 (Invivogen) for 3.5 hr. Inhibitors were prepared by serial dilution in DMSO. The media were removed and replaced with OptiMem containing inhibitors or 0.1% DMSO vehicle control. After 30 min incubation 5 μM Nigericin (Adipogen) was added for 1 hr. Cell free supernatants were removed and assessed by LDH assay (Roche) and ELISA for IL-1β according to the manufacturer’s instructions (mouse DuoSet, R&D Systems).

### Peritoneal Macrophage NLRP3 inflammasome activation assays

The generation of 129S6 human promoter-NLRP3 and 129S6 mouse promoter-NLRP3 mice been described previously(Koller et al., 2024; Snouwaert et al., 2016). Peritoneal macrophages were isolated from 3-month-old male 129S6-WT or 129S6-human promoter or mouse promoter *NLRP3* mice by lavage with 3-5 mL of PBS (no Ca^2+^ and no Mg^2+^). Cells from 2-4 individual mice were combined for each experiment. Fresh cells were then spun down and resuspended in complete growth media RPMI-1640 + GlutaMAX (Gibco)/ 10% fetal bovine serum (Corning)/ pen/strep (Caisson) and 50,000 - 100,000 cells (150µL) were then added to the appropriate wells of a 96-well TC plate. The cells were incubated overnight in a standard cell culture incubator (37°C; 5% CO_2_). The following day 50uL of GM-1 containing 400 ng/mL LPS (*E.coli* O26:B6; Sigma) was added to the appropriate wells (final concentration; 100 ng/mL LPS) and the cells were incubated for 5 hrs. The media was removed and replaced with Opti-Mem medium (without serum; Thermo) containing pre-determined dilutions of test compounds in replicate wells. After a 30 min incubation, 10 µM nigericin (final concentration; Sigma) or 4 mM ATP (final concentration; TOCRIS) in Opti-Mem medium with the corresponding concentration of test compound was added to the wells for an additional 1 hr. Positive control wells contain 10 µM nigericin or 4 mM ATP in Opti-Mem in the absence of test compound, while negative control wells contain Opti-Mem only. Supernatants are then transferred to a fresh 96-well plate for storage and assayed for IL-1β (mouse; DuoSet; R&D, DY401-05) and/or relative LDH levels (as a surrogate for pyroptosis) using a CytoTox 96 Kit (Promega; G1780).

### Sequence Alignments

Multiple protein sequence alignments were performed using CLUSTAL W Multiple Sequence Alignment Program (version 1.83). Protein sequences from the following GenBank accession numbers were used: Homo sapien (human) NLRP3 Q96P20.3, Macaca fascicularis (cynomolgus monkey; crab-eating macaque) NLRP3 XP_045246555, Chlorocebus sabaeus (African green monkey) NLRP3 XM_007990053.2, Mus musculus (mouse) NLRP3 NP_001346567.1, Rattus norvegicus (rat) NLRP3 NM_001191642.1, Oryctolagus cuniculus (European rabbit) NLRP3 QHZ00929.1 and Canis lupus familiaris (dog) NLRP3 [transcript variant X1] XM_038673023.1. For all species, the sequence alignments were restricted to the corresponding sequence of amino acids 131–694 in the human NACHT domain construct used in nanoDSF studies (Supp Fig. 4C).

### ADP-Glo™ Kinase Assay

To assess NLRP3 ATPase activity, the ADP-Glo™ Kinase Assay (Promega) was used according to the manufacturers supplied instructions. Specific assay conditions were as follows: 2.5 µM MBP-ΔNLRP3-HIS protein construct (CRELUX) (Hartman et al., 2024) was pre-incubated with pre-determined dilutions of test compounds for 15 min at 37°C prior to the addition of 25 µM of UltraPure ATP (Promega) for 2 hr at 37°C. The assay buffer was composed of 50 mM Tris, pH7.5, 150 mM NaCl, 10 mM MgCl2, 10% glycerol and 0.005% Tween-20), and replicate wells of each condition were tested in three separate experiments. Luminence was recorded with an integration time of 100 ms on either a Tecan Infinite® M1000 PRO or Molecular Devices SpectraMax® M5e.

### NanoDSF Assays

For the non-competitive thermal shift assay, 3 µM NLRP3^NACHT^ protein was incubated for 30 min on ice with 2% DMSO or 10 µM BAL-0028 or 10 µM MCC950 (1:3.3 ratio of protein to compound). For the competitive thermal shift assay, 3 µM NLRP3^NACHT^ protein was treated with 2% DMSO as the reference, with 20 µM MCC950 (1:6.7 ratio of protein to compound), or with 10 µM MCC950 and 10 µM BAL-0028 (1:6.7 ratio of protein to compounds) and incubated for 30 min on ice. The measurements were setup with a temperature ramp ranging from 20-90°C, a slope of 1.5°C·per minute and at 100% laser intensity. Data points are representative of three independent experiments.

### Drug Affinity Responsive Target Stability (DARTS) Assay

The DARTS assay was adapted from a published protocol(Pai et al., 2015). HEK293T cells (Merck) were cultured in DMEM + 10% FCS+ 50 U pen/strep + 1% Glutamax (all from Gibco). Cells were seeded at 3 x10^5^/ml in 10 cm^2^ dishes. The next day were transfected with 3-5 μg of plasmids, pEF6_human NLRP3-mCherry (a kind gift from Prof Kate Schroder, University of Queensland), pCDNA3.4 human NLRP3-Twin-Strep-tag or human NLRP3_1-688-Twin-Strep-tag (synthesized by GenScript) using the calcium phosphate method(Kwon and Firestein, 2013). Before cells were harvested 0.1-10 μM BAL-0028 or MCC950 or DMSO vehicle control was added. 24 hrs post transfection cells were harvested. Media were removed and cells were washed in PBS containing inhibitors or DMSO control, cells were lysed in lysis buffer (50 mM Tris-HCl pH 7.4, 150 mM NaCl, 1 mM ATP, 2 mM EDTA, 0.5% Igepal CA-630) containing protease inhibitors (complete mini protease inhibitor cocktail; Roche), benzonase and inhibitors or DMSO as indicated. Lysates were disrupted by passage through a 27-gauge needle and cleared by centrifugation at 14,000*g* for 10 min at 4 °C. Protein concentration was determined using a Pierce Rapid Gold BCA Protein Assay Kit (Thermo Fisher). Pronase and thermolysin 10 mg/mL, (Merck) were added at the indicated protease to protein ratio, for 15 min at room temperature. The reaction was stopped by addition of 20X protease inhibitor cocktail and incubated on ice for 10 min.

Protein samples were prepared with NuPAGE LDS sample buffer (Thermo Fisher) supplemented with 10 mM DTT. Samples were then resolved by SDS–PAGE using 4–20% Mini-PROTEAN TGX stain-free gels (Biorad) and transferred onto nitrocellulose membrane using the Trans-Blot Turbo transfer system (Biorad). Membranes were blocked in 5% (wt/vol) dried milk in TBS-T (10 mM Tris/HCl, pH 8, 150 mM NaCl and 0.05% (vol/vol) Tween-20) for 1 h at room temperature. Membranes were incubated with primary antibody NLRP3 clone D4D8T at 1:1,000 (15101, Cell Signaling Technology) diluted in 5% (wt/vol) dried milk in TBS-T and then with Peroxidase-AffiniPure Goat Anti-Rabbit IgG (Jackson ImmunoResearch) at 1 in 5,000. Membranes were developed using Clarity Western ECL substrate (Biorad) and then visualized using a Syngene G:Box.

### Mouse Plasma Protein Binding Determination

On the day of the experiment, pooled male CD-1 mouse EDTA-K^2^ plasma (BIOIVT) was thawed and centrifuged at 3220 rpm for 5 min and only plasma within range of pH 7.0 - 8.0 was used. Working solutions of BAL-0598 and the control compound Warfarin were prepared, and aliquots of working solutions were spiked into blank plasma matrix (10 – 99.5%) to achieve a final concentration of 2 µM, 10 µM, 30 µM or 100 µM of BAL-0598. The HT-Dialysis plate and the dialysis membranes (molecular weight cut off 12-14 KDa) were purchased from HT Dialysis LLC (Gales Ferry, CT). The dialysis buffer was composed of 100 mM sodium phosphate and 150 mM NaCl, pH 7.4 ± 0.1) and the stop solution was composed of acetonitrile containing 200 ng/mL tolbutamide and 200 ng/mL labetalol. Dialysis membranes were pretreated according to the manufacturer’s instructions and the dialysis instrument was prepared according to the manufacturer’s instructions. For T0 samples, aliquots of mouse plasma containing BAL-0598 or control compound were transferred in triplicate to the sample collection plate and mixed with blank buffer at 1:1 (v:v). Stop solution was then added and the plate was sealed and mixed at 800 rpm for 10 min and stored at 2-8°C until all samples were ready for analysis.

Ultracentrifugation was performed with a Beckman Coulter, OptimaTM L-90K. To prepare protein-free samples (F samples) for unbound determination, an aliquot of the pre-incubated matrix containing BAL-0598 or control compound was transferred to ultracentrifuge tubes (n=2) and subjected to ultracentrifugation at 37°C, 155000×g (35000rpm) for 4 hr. At the end of the ultracentrifugation, F sample aliquots were taken from the second layer down in the supernatant. To prepare the T4.5 samples, the residual aliquot of pre-incubated spiked plasma was placed into the incubator for 4 hr. These samples were then transferred to new 96 well plates and an equal volume of buffer or plasma was added to each well. Stop solution was then added to these samples and the mixture was vortexed and centrifuged at 4000 rpm for about 20 minutes. After dialysis or ultracentrifugation, compound levels were determined by LC-MS/MS analysis. Dialysis: %Unbound = 100 × F / T; %Bound = 100 - %Unbound; %Recovery = 100 × (F + T) / T0; F = Free compound concentration as determined by the calculated concentration on the buffer side of the membrane; T = Total compound concentration as determined by the calculated concentration on the matrix side of the membrane; T0 = Total compound concentration as determined by the calculated concentration in matrix before dialysis. Ultracentrifugation: %Unbound = 100 × F / Mean of T4.5; %Bound = 100 - %Unbound; %Remaining = 100 × Mean T4.5 / Mean of T0.

### In vivo LPS/ATP-Induced Peritonitis Mouse Model

All experiments were approved by the Institutional Animal Care and Use Committee (IACUC) of BioAge Labs (Richmond, CA). Female, 4-mo old 129S6-hNLRP3 mice were group housed during acclimation before experiment and randomized to different treatment group cages on the day of experiment. Mice were housed conventionally in a constant temperature (18–26°C) and humidity (30%–70%) animal room with a 12/12 h light/dark cycle and free access to food (5LG4 - LabDiet® JL Rat and Mouse /Irr 6F) and water (M-WB-300A, Innovive) ad libitum. Temperature and relative humidity were monitored daily. The animals were identified by cage card and tail mark during the study to identify the animal individually.

In the peritonitis studies, the mice were intraperitoneally injected with LPS at 40 µg/kg, followed two hours later by an intraperitoneal injection of ATP disodium salt at 304.4 mg/kg. Animals not treated with ATP (i.e. Group 1) were dosed with the appropriate volume of PBS. BAL-0598 was administered to fed mice once at 3, 10, 30 or 100 mg/kg by oral gavage (PO) at 10 µL/g following facility SOPs. Vehicle (20% propylene glycol + 40% PEG-400 + 10% ethanol) only or BAL-0598 was PO dosed one hour prior to ATP IP administration which approximately corresponded to the plasma T_max_ of BAL-0598 (data not shown) at the time of ATP administration. Thirty min after ATP administration, submental bleeds were collected to blood collection tube with EDTA-K^2^ (BD 365974). Blood samples were centrifuged at approximately 4°C, 13,000 x g for 10 min to collect plasma to determine total plasma BAL-0598 concentrations by LC-MS/MS (Quintara Discovery, Hayward, CA), and then the mice were immediately euthanized with isoflurane and the peritoneal cavity was lavaged with 3 mL of PBS to collect the PLF. Plasma BAL-0598 levels were determined by LC-MS/MS. The level of mature IL-1β and IL-6 in the PLF was determined by ELISA (R&D Systems).

### In Vivo IC_50_ Determination

For in vivo IC_50_ determinations, the absolute values of IL-1β in the PLF were plotted against the unbound BAL-0598 concentrations (nM) determined at the time of euthanasia (i.e. 1.5 hr after BAL-0598 or vehicle dosing). Unbound BAL-0598 values were calculated from the total BAL-0598 measured in the plasma of individual mice using an unbound value of 1.61% (Supp. Table 1). Relative IC_50_ values (i.e., the unbound BAL-0598 concentration required to bring the curve down to a point halfway between the “Top” and “Bottom” plateaus of the curve) were calculated by non-linear regression analysis using GraphPad PRISM version 10.1.1. The upper plateaus were anchored using the IL-1b values derived from the vehicle dosed mouse promoter 129S6-hNLRP3 mice treated with LPS + ATP with the unbound BAL-0598 set to a default value of 0.002 nM (one order of magnitude below the LLOQ).

### U937 expressing NLRP3-AID associated mutant assays

NLRP3-deficient U937 cells reconstituted with doxycycline-inducible NLRP3 mutants have been previously described(Cosson et al. 2024). 0.3×10^6^ U937 cells/ml were platted in Roswell Park Memorial Institute (RPMI) 1640 GlutaMax^TM^-I supplemented with 1X PS and 10% FBS (Gibco). The next day, cells were treated the BAL-0028 or MCC950 (1000, 200, 40 nM, final DMSO concentration 0.05% in all conditions). 15 min later, cells were treated with doxycycline (1 μg/ml, 3h, Sigma), LPS (40 ng/ml, 2h, O111:B4, Sigma) and nigericin (15 μg/ml, Invivogen) before time-lapse imaging using CQ1 high content screening microscope (Yokogawa). PI (1.25 μg/ml) and Hoechst (0.2 μg/ml) were added 2h before imaging. 2 images/well were taken every 15 min for 2h using 10X objectives (UPLSAPO 10X/0.4). Image analysis was performed as previously described(Cosson et al., 2024).

### Data Analyses

The absolute values of IL-1β in the cell culture supernatants were normalized relative to the mean level of IL-1β present in the positive control (i.e., the maximum response) and shown as a percent of maximum response. These values were calculated independently within each experiment. Relative IC_50_ values (i.e., the BAL-0028 or MCC950 concentration required to bring the curve down to a point halfway between the “Top” and “Bottom” plateaus of the curve) were calculated by non-linear regression analysis using GraphPad PRISM version 9.5.0.

For cytotoxicity (LDH release) calculations, raw absorbance values at 490 nm (OD_490_) of cell culture supernatants were normalized relative to the OD_490_ of the maximal LDH release control (10X Lysis Reagent). Percent cytotoxicity was calculated using the formula, %Cytotoxicity = 100 X (Experimental LDH release (OD_490_) / Maximal LDH release (OD_490_)).

### Data Availability

All raw data are available upon request.

## ACKNOWLEDGEMENTS

We thank Prof Kate Schroder (University of Queensland, Australia) for the pEF6 human NLRP3-mCherry construct and Dr. Joana Sá-Pessoa (Queen’s University Belfast, UK) for guidance with the immortalization of BMDM.

## AUTHOR CONTRIBUTIONS

K. Wilhelmsen, A. Deshpande, S. Tronnes, M. Mahanta, M. Banicki, M. Cochran, S. Cowdin, C. Portillo, S. Yan, A. Duisembekova, R. Riou, M. Marleaux, I.V. Hochheiser, H. Buthmann, W. Wang, M. Cranston, C. McKee, T. Mawhinney, E. McKay performed experiments. B. Morgan contributed to the development of BAL-0028. K. Wilhelmsen, A. Deshpande, S. Tronnes, G. Hartman, R. Hughes, Y. Wang, B.F. Py, M. Geyer, R.C. Coll conceptualized the studies. K. Wilhelmsen and R.C. Coll wrote the manuscript, which was modified after input from all authors. K. Fortney, R. Montgomery and P. Rubin acquired funding and provided supervision.

## FIGURE LEGENDS

**Fig S1. (related to Figure 1).**
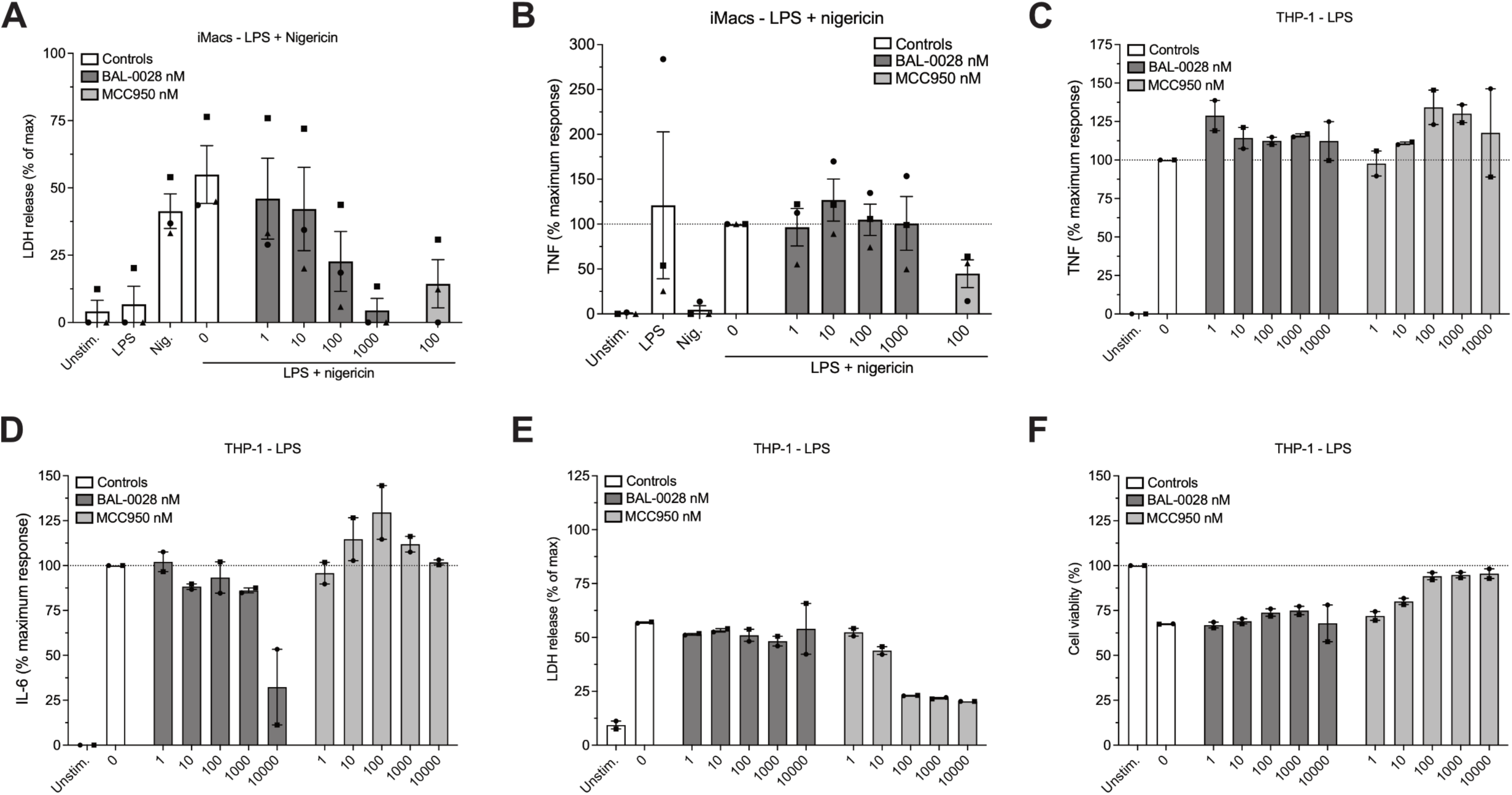
(A-B) Effect of BAL-0028 and MCC950 on LDH release (A) and TNF release (B) from LPS and nigericin stimulated iMacs. (C, D) Comparison of BAL-0028 and MCC950 pretreatment in PMA-differentiated THP-1s on LPS-induced secretion of (C) TNF and (D) IL-6. (E-F) Effect of BAL-0028 and MCC950 treatment in PMA-differentiated THP-1s on (E) cytotoxicity (LDH release) and (F) and cell viability (CellTiter-Blue assay). (A-F) Bar chart symbols show average values relative to vehicle control from independent experiments (indicated by different symbols) performed in triplicate+/− S.E.M. (A-B) N=3. (C-F) N=2.

**Fig S2. (related to Figure 2).**
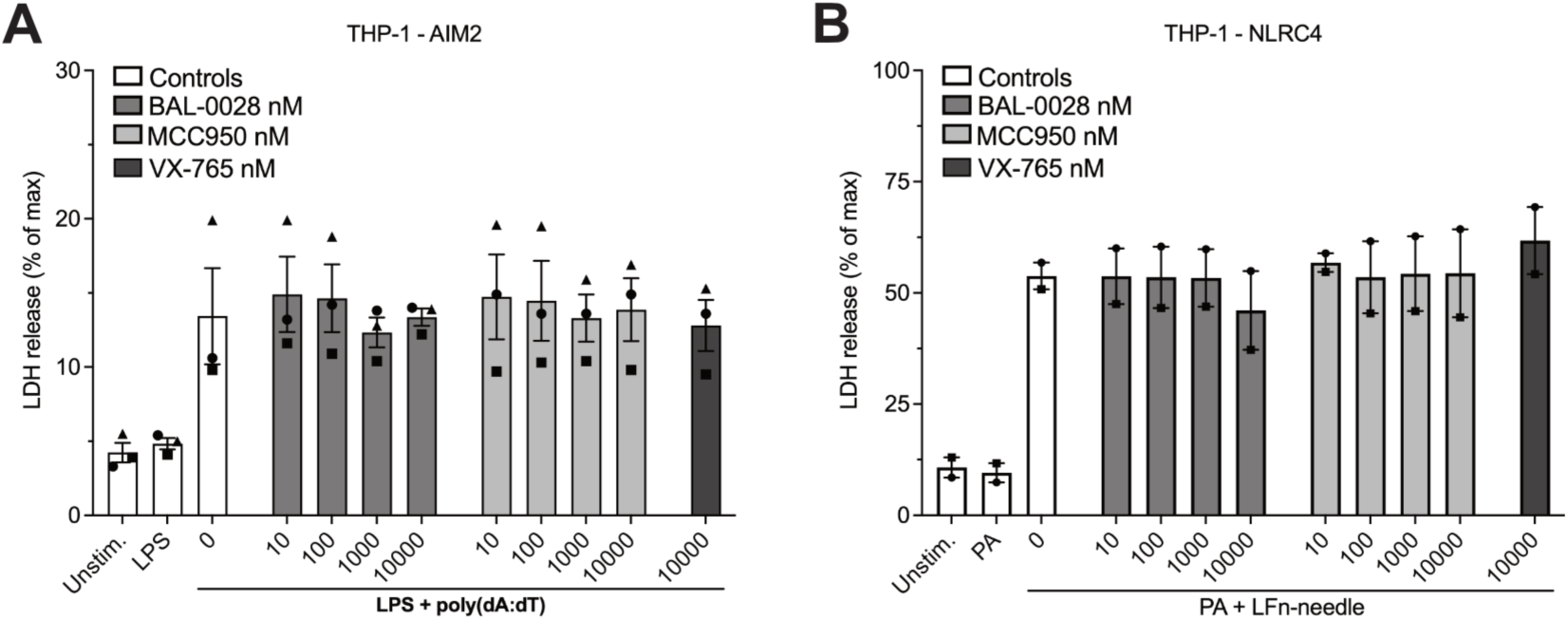
Effects of BAL-0028, MCC950 and VX-765 on LDH release from PMA differentiated THP-1 cells stimulated with (A) LPS and transfected with poly(dA:dT) or (B) protective antigen (PA) and Lfn-needle protein. Graph symbols show average values from independent experiments performed in triplicate (indicated by different symbols) +/− S.E.M. (A) N=3 and (B) N=2.

**Fig S3. (related to Figure 3).**
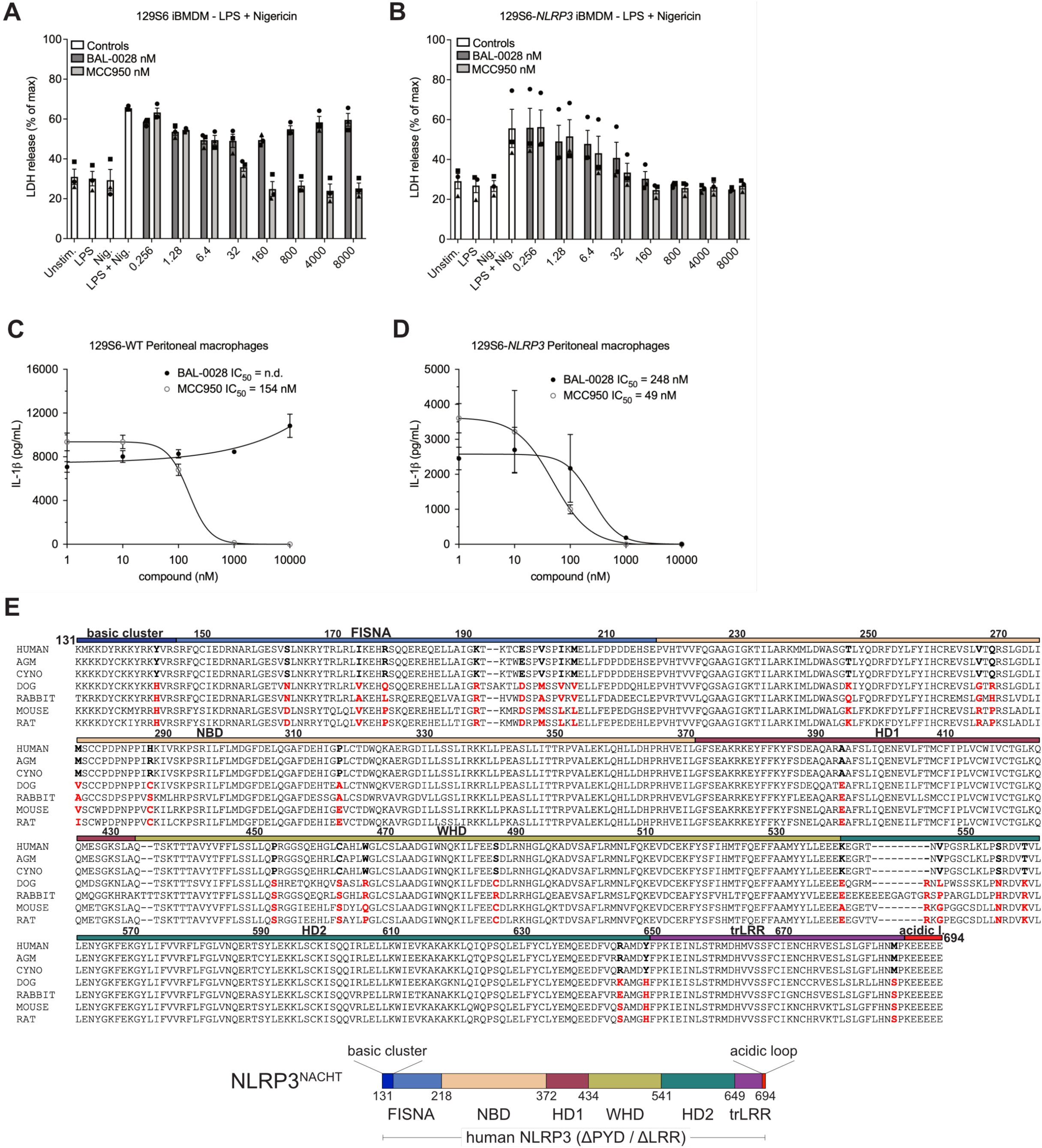
(A-B) Comparison of BAL-0028 and MCC950 in LDH release assays from (A) wild-type 129S6 iBMDM and (B) 129S6-human promoter *NLRP3* iBMDM cells stimulated with LPS and nigericin. Graph symbols show average LDH values +/− S.E.M. from N=3 independent experiments (indicated by different symbols) performed in triplicate. (C-D) Comparison of BAL-0028 and MCC950 in IL-1β release assays from (C) wild-type 129S6 and (D) 129S6-human promoter *NLRP3* primary peritoneal macrophages stimulated with LPS and nigericin. Graph symbols show average IL-1β values relative to vehicle control +/−S.D. from one experiment performed in duplicate. IC50 curves fitted by non-linear regression analysis. (E) Multiple sequence alignment of human, African green monkey (AGM), Cynomolgus monkey (CYNO), dog, rabbit, mouse, and rat NLRP3 protein sequences restricted to the corresponding sequence of amino acids 131 – 694 in the human NACHT domain construct shown below the alignment.

**Fig. S4 (related to Figure 4).**
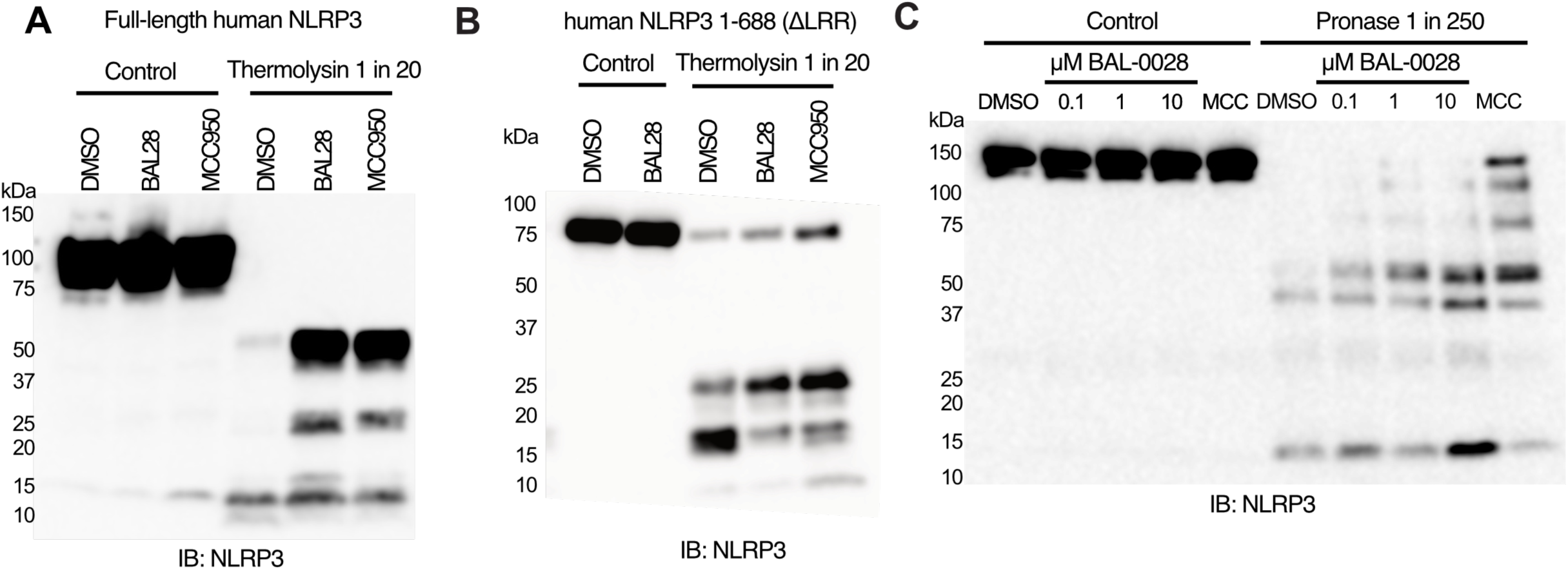
(A-C) Western blots showing NLRP3 expression and degradation in DARTS assays performed with (A) full length human NLRP3-Twin-Strep-tag (B) human NLRP3 1-688-ΔLRR-Twin-Strep-tag or (C) human-NLRP3-mCherry. Cells and cell lysates were treated with 10 μM BAL-0028 or MCC950 or DMSO control (A-B) or 0.1-10 μM BAL-0028, 10 μM MCC950 or DMSO control (C). Blots shown are representative of (A) N=3, (B) N=2) and (C) N=4 independent experiments.

**Fig. S5 (related to Figure 5).**
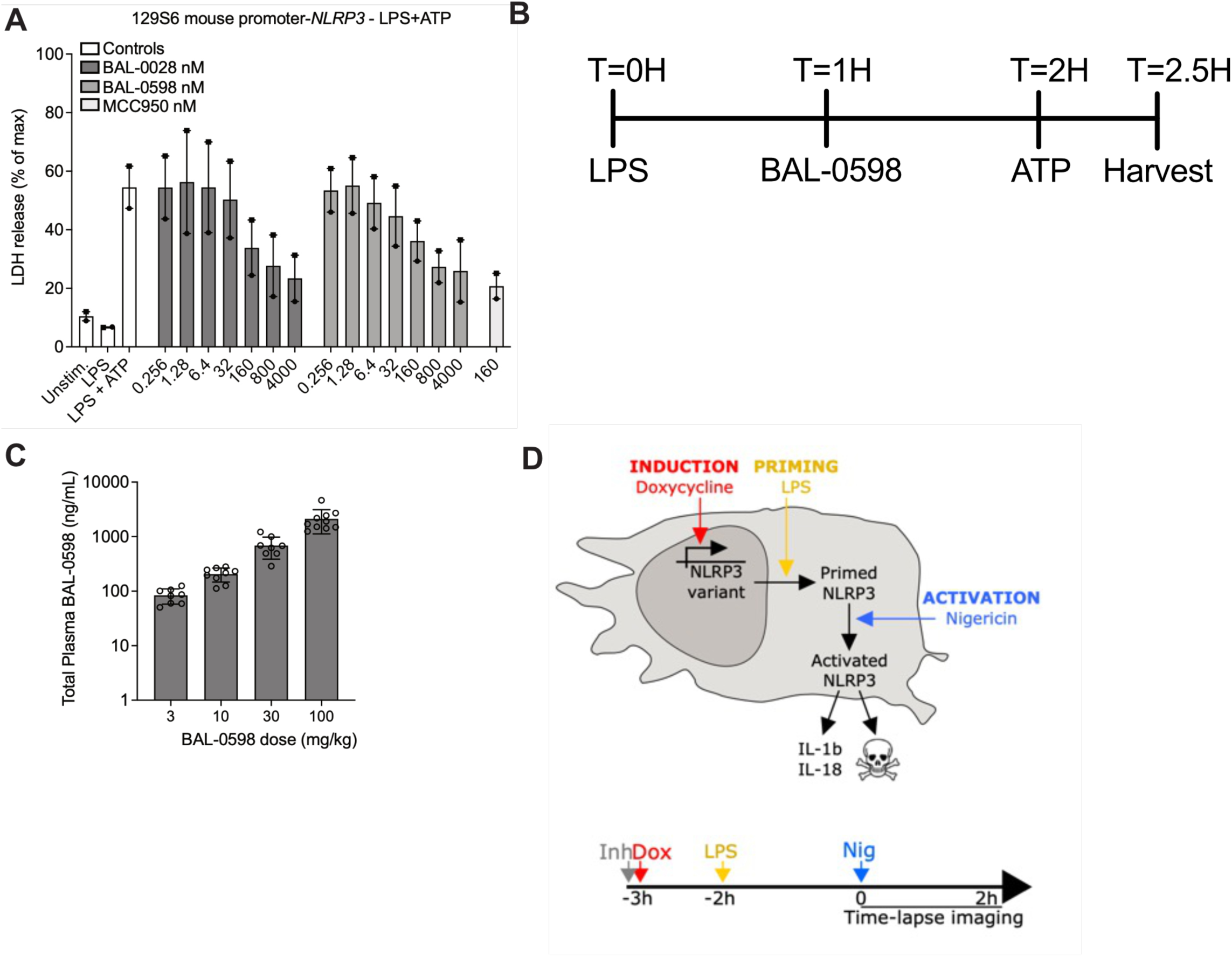
(A) Comparison of BAL-0598, BAL-0028 and MCC950 in an LDH release assay in primary peritoneal macrophages isolated from 129S6 mouse promoter-*NLRP3* mice stimulated with LPS and ATP. Graph symbols show average LDH values +/− S.E.M. from N=2 independent experiments performed in duplicate. (B) Schematic illustration of peritonitis model and BAL-0598 dosing. (C) Total plasma BAL-0598 levels from BAL-0598 oral gavage treated mice. Graph symbols show values from individual mice +/− S.D. (D) Schematic illustration of U937 NLRP3 and NLRP3-AID mutant cell model.

**Supplemental Table 1.**
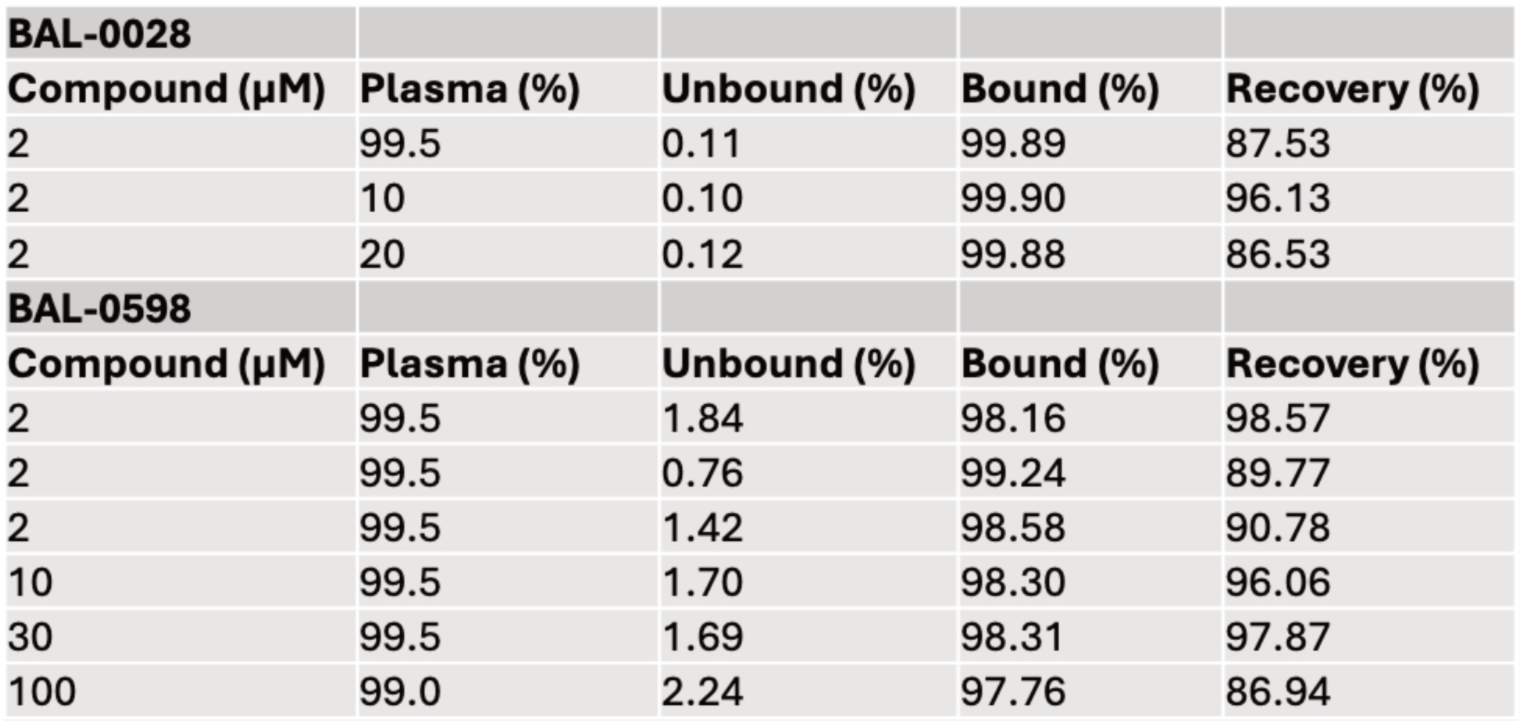
Plasma protein binding of BAL-0028 and BAL-0598.

